# Induction of a regulatory myeloid program in bacterial sepsis and severe COVID-19

**DOI:** 10.1101/2020.09.02.280180

**Authors:** Miguel Reyes, Michael R. Filbin, Roby P. Bhattacharyya, Abraham Sonny, Arnav Mehta, Kianna Billman, Kyle R. Kays, Mayra Pinilla-Vera, Maura E. Benson, MGH COVID-19 Collection & Processing Team, Lisa A. Cosimi, Deborah T. Hung, Bruce D. Levy, Alexandra-Chloe Villani, Moshe Sade-Feldman, Rebecca M. Baron, Marcia B. Goldberg, Paul C. Blainey, Nir Hacohen

**Affiliations:** Broad Institute of MIT and Harvard, Cambridge, MA, USA; Department of Biological Engineering, Massachusetts Institute of Technology, Cambridge, MA, USA; Department of Emergency Medicine, Massachusetts General Hospital and Harvard Medical School, Boston, MA, USA; Center for Bacterial Pathogenesis, Division of Infectious Diseases, Department of Medicine, Massachusetts General Hospital and Harvard Medical School, Boston, MA, USA; Department of Anesthesia, Critical Care & Pain Medicine, Massachusetts General Hospital and Harvard Medical School, Boston, MA, USA; Center for Cancer Research, Massachusetts General Hospital and Harvard Medical School, Boston, MA, USA; Division of Pulmonary and Critical Care Medicine, Department of Medicine, Brigham and Women’s Hospital and Harvard Medical School, Boston, MA, USA; Center for Immunology and Inflammatory Diseases, Massachusetts General Hospital and Harvard Medical School, Boston, MA, USA

## Abstract

A recent estimate suggests that one in five deaths globally are associated with sepsis^1^. To date, no targeted treatment is available for this syndrome, likely due to substantial patient heterogeneity^2,3^ and our lack of insight into sepsis immunopathology^4^. These issues are highlighted by the current COVID-19 pandemic, wherein many clinical manifestations of severe SARS-CoV-2 infection parallel bacterial sepsis^5–8^. We previously reported an expanded CD14+ monocyte state, MS1, in patients with bacterial sepsis or non-infectious critical illness, and validated its expansion in sepsis across thousands of patients using public transcriptomic data^9^. Despite its marked expansion in the circulation of bacterial sepsis patients, its relevance to viral sepsis and association with disease outcomes have not been examined. In addition, the ontogeny and function of this monocyte state remain poorly characterized. Using public transcriptomic data, we show that the expression of the MS1 program is associated with sepsis mortality and is up-regulated in monocytes from patients with severe COVID-19. We found that blood plasma from bacterial sepsis or COVID-19 patients with severe disease induces emergency myelopoiesis and expression of the MS1 program, which are dependent on the cytokines IL-6 and IL-10. Finally, we demonstrate that MS1 cells are broadly immunosuppressive, similar to monocytic myeloid-derived suppressor cells (MDSCs), and have decreased responsiveness to stimulation. Our findings highlight the utility of regulatory myeloid cells in sepsis prognosis, and the role of systemic cytokines in inducing emergency myelopoiesis during severe bacterial and SARS-CoV-2 infections.

## Main Text

Sepsis is associated with profound alterations in the peripheral immune compartment, including marked reduction in lymphocyte counts^10–12^ and phenotypic alteration of myeloid cells^13–15^. Monocytes from sepsis patients have decreased responsiveness to stimuli^15–17^ and have lower expression of HLA-DR^18–23^ characteristic of monocytic MDSCs. In line with these findings, we recently reported an expanded CD14+ monocyte state in sepsis patients, MS1, reminiscent of monocytic MDSCs^9^. Previous studies have reported opposing effects of MDSCs on sepsis outcomes^24–27^, warranting further investigation into their function and association with patient prognosis. Signatures similar to MS1 have also been described recently in severe SARS-CoV-2 infections^28,29^, but have not been systematically analyzed across multiple cohorts. Here, we identify the gene expression programs associated with the MS1 cell state and determine their relationship with disease severity in both bacterial sepsis and COVID-19. We identify circulating host factors that result in the induction of the MS1 gene module, and characterize the response of MS1 cells to stimulation and their effects on other cell types.

To characterize the gene expression program associated with the MS1 cell state, we analyzed existing monocyte scRNA-seq data from sepsis patients and controls^9^ using consensus non-negative matrix factorization^30^ (cNMF; **Figure 1a, Supplementary Fig. 1a-d**). We found a gene expression program that includes the MS1 marker genes *RETN, ALOX5AP*, and *IL1R2* among the top 20 genes with highest loadings (**Supplementary Fig. 1d**), has higher usage in sepsis patients (**Supplementary Fig. 1e**, FDR < 0.001), and correlates with the fractional abundance of MS1 cells in each patient (**Supplementary Fig. 1f**, r = 0.73, p < 0.001). This module negatively correlates with the usage of a class II major histocompatibility complex module (MHC-II module, **Supplementary Fig. 1g**), consistent with our observation that MS1 cells have lower surface expression of HLA-DR^9^. Co-expression analysis within the MS1 module reveals that most genes are highly correlated with *S100A8* (**Figure 1b**), suggesting its role as a core driver of the MS1 gene expression program. This gene and its partner *S100A9* have been implicated in the development of MDSCs in cancer^31^ and sepsis^24,32^, indicating a similarity between monocytic MDSCs and the MS1 cell state.

**Figure 1.**
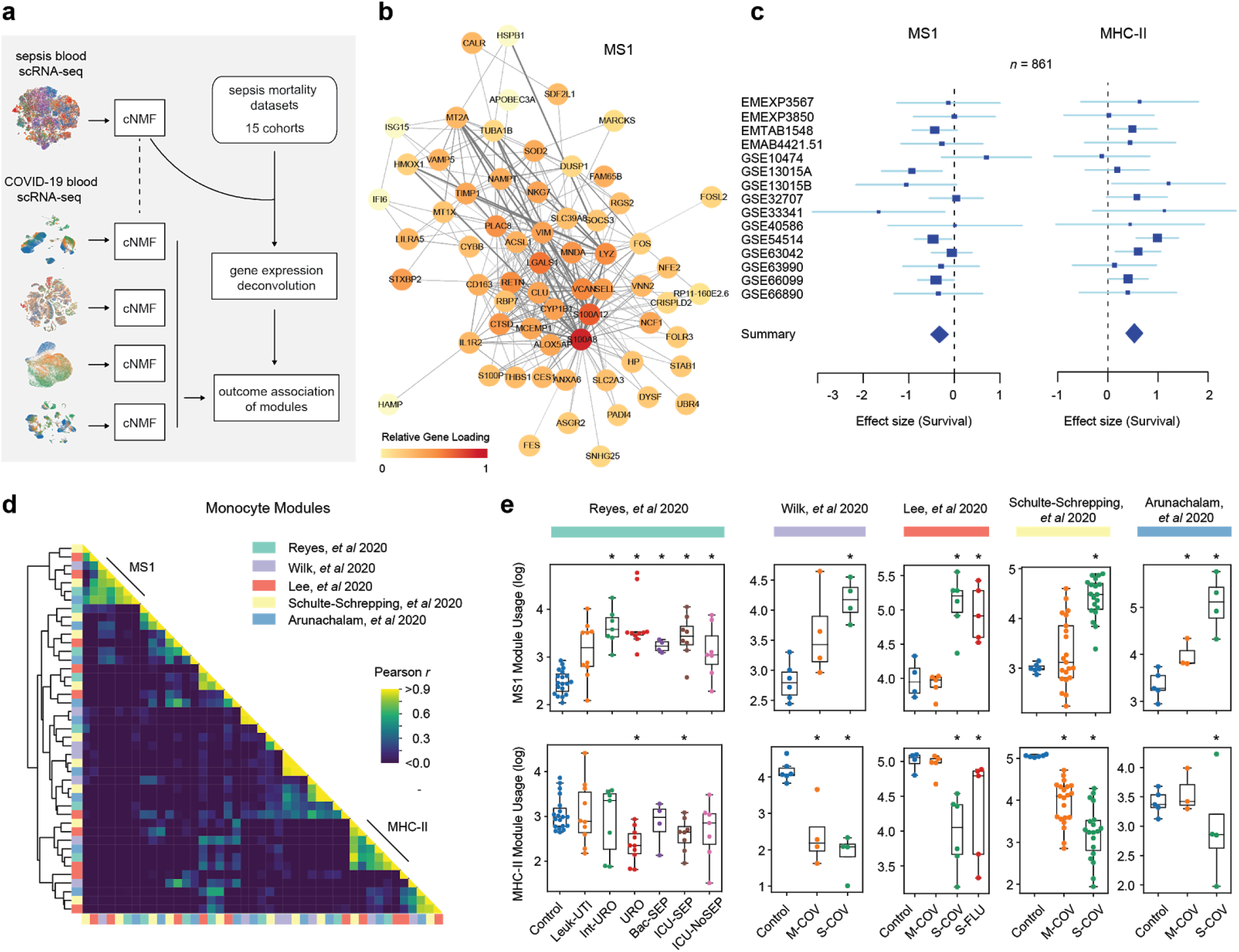
The MS1 gene expression program is associated with disease severity in bacterial sepsis and severe COVID-19. (a) Schematic of the analysis. (b) Correlation network of the MS1 module. Edge thickness is proportional to the correlation value between each pair of genes. Node colors are proportional to the expression level in log(TPM) of each gene in the module. (c) Forest plots showing the effect size on survival (log2 standardized mean difference) of inferred MS1 (left) or MHC-II (right) module usage in each dataset from bulk gene expression deconvolution. Accession numbers of the data from each study are listed on the left. Boxes indicate the effect size in an individual study, with whiskers extending to the 95% confidence interval. Size of the box is proportional to the relative sample size of the study. Diamonds represent the summary effect size among the patient groups, determined by integrating the standardized mean differences across all studies. The width of the diamond corresponds to its 95% confidence interval. (d) Correlation matrix of the gene weights (z-scored) for the monocyte gene expression modules across the 4 datasets. Gene modules were unbiasedly derived from each dataset using consensus non-negative matrix factorization. (e) Mean usage (log) of the MS1 (top) and MHC-II (bottom) modules in monocytes for each patient across patient groups for each dataset. Asterisks indicate an FDR < 0.05, computed by comparing each disease state with healthy controls (two-tailed Wilcoxon rank-sum test, corrected for testing of multiple modules). Boxes show the median and IQR for each patient cohort, with whiskers extending to 1.5 IQR in either direction from the top or bottom quartile. Detailed description of the patient cohorts and number of cells and patients for each dataset in (d,e) are outlined in the corresponding publications^7,9,28,34,35^. Control, healthy controls; Leuk-UTI, urinary tract infection with leukocytosis; Int-URO, intermediate urosepsis; URO, urosepsis; Bac-SEP, sepsis with confirmed bacteremia; ICU-SEP, intensive care with sepsis; ICU-NoSEP, intensive care without sepsis; M-COV, mild COVID-19; S-COV, severe COVID-19; S-FLU, severe influenza A.

Given its marked expansion, we hypothesized that expression of the MS1 program may be associated with worse outcome in bacterial sepsis. Using the transcriptional data from our earlier study^9^, the usage of the MS1 program correlates with sepsis severity in ICU patients but not in patients presenting to the emergency department with milder disease (**Supplementary Fig. 1h**). Using gene expression deconvolution, we estimated the usage of the monocyte gene expression modules in 15 datasets included in a recent meta-analysis examining sepsis mortality^33^ (**Methods, Supplementary Figure 1c, Supplementary Table 1**). We found that expression of the MS1 program is negatively associated with patient survival (effect size = -0.32, FDR < 0.01), whereas MHC-II module usage has the opposite effect (effect size = 0.53, FDR < 0.01; **Figure 1c**). These results show that expansion of MS1 cells is associated with worse infection outcomes, suggesting its prognostic value for sepsis patients.

To determine whether MS1 cells are similarly expanded in severe COVID-19, we analyzed four COVID-19 scRNA-seq datasets^7,28,34,35^ independently and identified gene expression modules in CD14+ monocytes from each dataset using an unbiased cNMF method (**Figure 1a, Supplementary Figure 2**). In each of the four studies, we found gene expression modules corresponding to the MS1 and MHC-II modules, as evidenced by their strong correlation (Pearson *r* > 0.8) with modules from our sepsis data (**Figure 1d**). Similar to the trends we observed in bacterial sepsis, CD14+ monocytes from patients with severe SARS-CoV-2 or influenza A infections have significantly higher and lower usage of the MS1 and MHC-II modules, respectively (FDR < 0.05; **Figure 1e**), and have increased MS1 scores compared with controls (p < 0.01; **Supplementary Figure 2**). These findings show that the MS1 cell state is expanded in both bacterial and viral sepsis syndromes, suggesting its utility as a signature for infection severity regardless of etiology.

We previously demonstrated that MS1-like cells can be derived from immature progenitors through stimulation of total bone marrow mononuclear cells (BMMCs) with LPS or Pam3CSK4^9^. Because BMMCs contain a heterogeneous mix of hematopoietic stem and progenitor cells (HSPCs) and mature immune cells, potential paracrine interactions among the cell types confound identification of the precise factors that cause MS1 induction. Given this limitation, we sought a method for inducing the MS1 program directly from CD34+ HSPCs purified from bone marrow. We hypothesized that cytokines circulating in the blood of sepsis patients might induce the differentiation of MS1 cells directly from HSPCs. Upon culturing HSPCs isolated from healthy human bone marrow with plasma from urosepsis (URO) patients or healthy controls (Control) (**Figure 2a**), we found that sepsis plasma significantly stimulated the production of monocytes and neutrophils compared with healthy plasma (**Figure 2b**; p = 0.025 and 0.004 for CD34-CD11b+CD14+ and CD34-CD11b+CD15+ cells, respectively). Single cell analysis of the differentiated cell populations showed clear trajectories of myeloid differentiation (**Figure 2d, Supplementary Fig. 3a-b, Methods**); importantly, we observed that incubation of HSPCs in plasma from URO patients resulted in the emergence of CD14+ cells with high MS1 scores compared with Control (**Figure 2d;** p < 0.01). cNMF analysis of the scRNA-seq data identified modules similar to the MS1 and MHC-II modules derived from patient PBMC data, as evidenced by strong correlations between their gene loadings (Pearson *r* = 0.73 and 0.78, respectively; **Figure 2f, Supplementary Fig. 3c**). The MS1 and MHC-II modules are also significantly up- or down-regulated (p < 0.01), respectively, in CD14+ cells derived from HSPCs incubated with sepsis plasma (**Figure 2f**). These data support our hypothesis that cytokines circulating in the blood of sepsis patients can induce the differentiation of MS1 cells.

**Figure 2.**
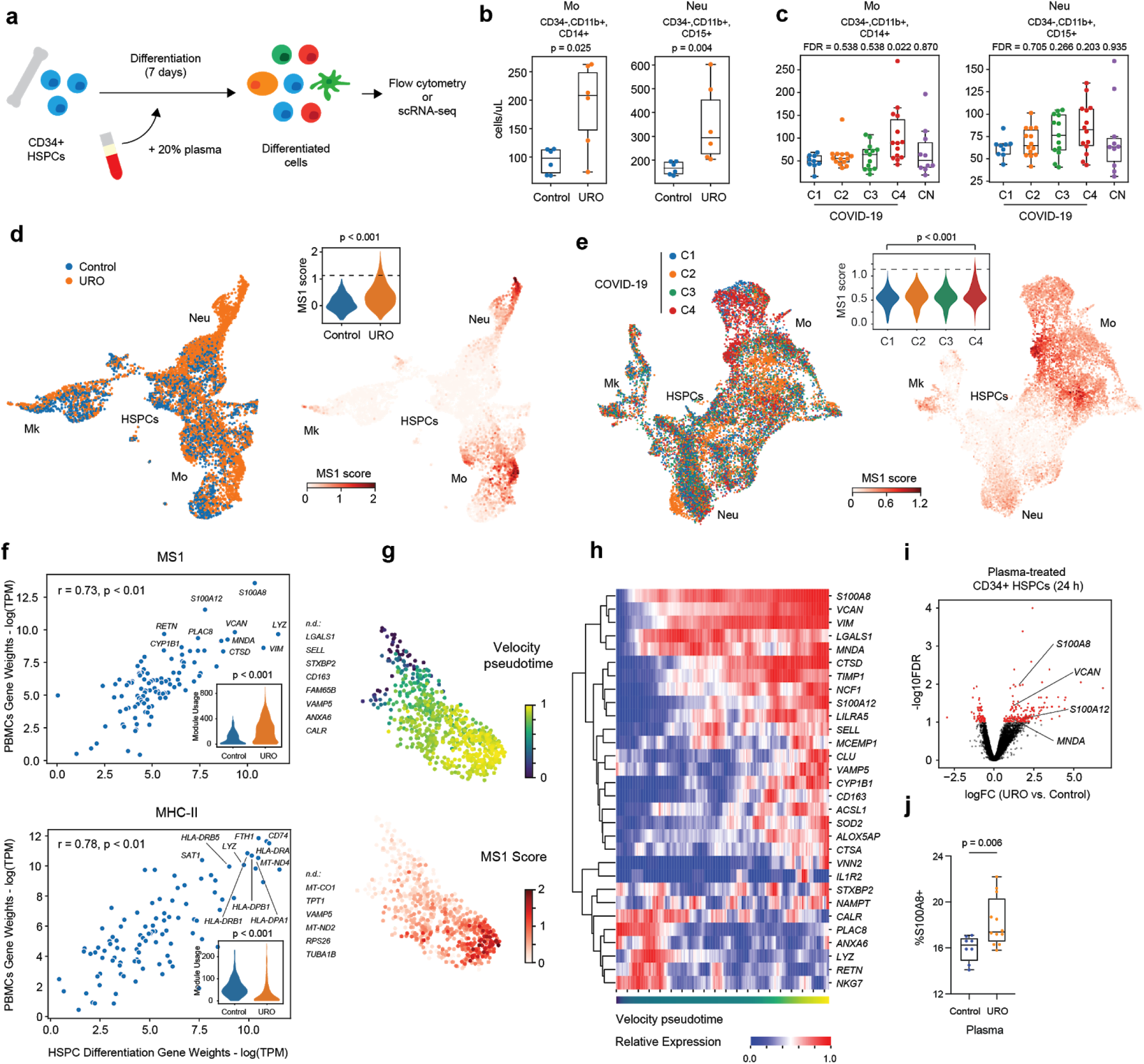
Sepsis and COVID-19 plasma induce the MS1 program from HSPCs. (a) Schematic of the plasma-incubation experiment. (b-c) Number of CD34-, CD11b+, CD14+ (left) and CD34-, CD11b+, CD15+ (right) produced after incubation of CD34+ HSPCs in medium with Control or indicated patient plasma for 7 days. Experiments in (b) were performed with *n* = 6 for each condition (3 plasma donors with 2 technical replicates). P-values are calculated using a two-tailed Wilcoxon rank-sum test. Experiments in (c) were performed with *n =* 9, 14, 14, 14, and 10 donors for C1-C4, and CN, respectively (**Methods**). FDR values are shown when comparing each disease state with C1 (two-tailed Wilcoxon rank-sum test, corrected for testing of multiple cohorts). Experiments in (b) and (c) were repeated in an independent bone marrow donor with similar results. (d-e) UMAP projections of scRNA-seq data from the experiment depicted in (a). Colors indicate which plasma pool the CD34+ progenitors were treated with (left) or the MS1 score (**Methods**) of each cell (right). Major immune cell types are labeled based on expression of known marker genes. The experiment in (d) was performed on 2 healthy bone marrow donors with 2 plasma donors for each condition; a total of 3,039 and 5,254 cells were profiled for Control and URO plasma treatment, respectively. The experiment in (e) was performed on 2 healthy bone marrow donors with pooled plasma from all donors in (c); a total of 4,449, 4,591, 3,129 and 3,711 cells were profiled for C1-C4 plasma, respectively. Inset shows a violin plot of MS1 scores for CD14-expressing cells from each condition. Dashed line indicates MS1 score in MS1 cells from the PBMC dataset. (f) Gene weight correlation between the MS1 (top) or MHC-II (bottom) modules detected in the plasma-incubation (x-axis) and patient PBMC datasets (y-axis). Significance of the correlations (Pearson *r*) are calculated with a permutation test. Genes which are not detected (n.d.) in the plasma incubation module but are among the top 30 for the corresponding module in the PBMC dataset are indicated. Insets show a violin plot of module usage across different plasma treatment conditions. (g) UMAP projection of Leiden cluster 6 (MS1 cells differentiating from CD34+ progenitors). Cells are colored by relative velocity pseudotime (top) and MS1 score (bottom). (h) Clustered heatmap of the top 30 genes in the MS1 module. Columns indicate individual cells ordered by velocity pseudotime. Expression values are z-score normalized for each gene. (i) Volcano plot showing differential expression analysis results (exact test) between CD34+ HSPCs treated with Control and URO plasma for 24 h. Genes with FDR < 0.1 are highlighted in red, and selected genes that are up-regulated early in the MS1 trajectory are labeled. *n* = 4 experiments were performed for each condition (2 plasma donors with 2 technical replicates). (j) S100A8 intracellular staining of CD34+ HSPCs treated with Control and URO plasma for 24 h. *n* = 4 and 6 plasma donors for Control and URO, respectively, with 2 technical replicates each. Mk, megakaryocytes; Neu, neutrophils; Mo, monocytes; HSPC, hematopoietic stem and progenitor cells; TPM, transcripts-per-million.

Due to the expansion of MS1 in severe COVID-19, we hypothesized that similar effects may be observed when incubating HSPCs with plasma from severe COVID-19 patients. We performed the same experiments with heat-inactivated plasma from SARS-CoV-2 infected patients with varied disease severity (C1-C4) and uninfected controls (CN; **Methods, Supplementary Fig. 3e-f, Supplementary Table 2**). Indeed, plasma from patients who eventually died of COVID-19 (C4) stimulated the production of monocytes more strongly than non-hospitalized COVID-19 patients (C1, FDR = 0.02; **Figure 2c**). Compared with C1 plasma, C4 plasma induced the production of CD14+ cells with higher MS1 scores (FDR < 0.01; **Figure 2e**) and caused increased and decreased usage of the MS1 and MHC-II modules, respectively (p < 0.01; **Supplementary Fig. 3g-h**). These findings further highlight the similarities between bacterial sepsis and severe COVID-19, and support our hypothesis that circulating factors from patients with severe disease stimulate the induction of MS1 cells.

Interestingly, upon incubation of HSPCs with URO plasma, we observed cells with high MS1 scores and a module of genes similar to MS1 in the emerging neutrophil population (**Figure 2d, Supplementary Fig. 3c**). Moreover, neutrophils sorted from critically-ill patients (ICU) and bacteremic (BAC) patients express a module which includes *S100A8* among the genes with highest loadings (**Supplementary Fig. 4a-c**) and correlates with the MS1-like module in neutrophils generated from incubating HSPCs with URO plasma (Pearson *r* = 0.58, **Supplementary Fig. 4d**). MS1 marker genes are also among the top differentially expressed genes in neutrophils between healthy control subjects and ICU or BAC patients (**Supplementary Fig. 4e**). These findings are consistent with previous reports of MDSCs having both granulocytic and monocytic subtypes^36^, and further highlight the similarity between the MS1 cell state and MDSCs.

Analysis of the monocyte differentiation trajectory shows different pseudo-temporal dynamics of the MS1 genes (**Figure 2g**). Of note, a number of genes, including *S100A8, VCAN*, and *MNDA*, are expressed early and remain up-regulated during the differentiation trajectory (**Figure 2h)**. Short term stimulation (24 h) of CD34+ HSPCs with sepsis plasma results in the up-regulation of these genes (**Figure 2i, Supplementary Table 3**) and an increase in the fraction of S100A8+ cells in the CD34+ population (**Figure 2j**). These genes are among the core genes in the MS1 module (**Figure 1b**), and their up-regulation early is in line with our hypothesis that *S100A8* is an important factor driving the induction and differentiation of MS1 cells from HSPCs.

To determine which circulating factors induce the increased production of myeloid cells from HSPCs, we analyzed the levels of inflammatory cytokines implicated in cytokine storms^37^ in the plasma of COVID-19 patients (**Supplementary Figure 5**). We found that IL-6 strongly correlated with higher production of CD14+ cells from HSPCs (**Figure 3a**), suggesting its involvement in the induction of emergency myelopoiesis. To test this hypothesis, we added plasma from urosepsis or COVID-19 patients, or corresponding controls, to IL6R-knockout or control CD34+ HSPCs and observed a marked reduction in CD14+ cell production across all conditions for IL6R-knockout CD34+ HSPCs (**Figure 3b**). We also observed a reduction, albeit weaker, specifically for URO and C4 plasma when wild-type HSPCs are differentiated in the presence of IL-6 neutralizing antibodies (**Figure 3c**). These findings highlight the role of circulating IL-6 in inducing emergency myelopoiesis during severe infections.

**Figure 3.**
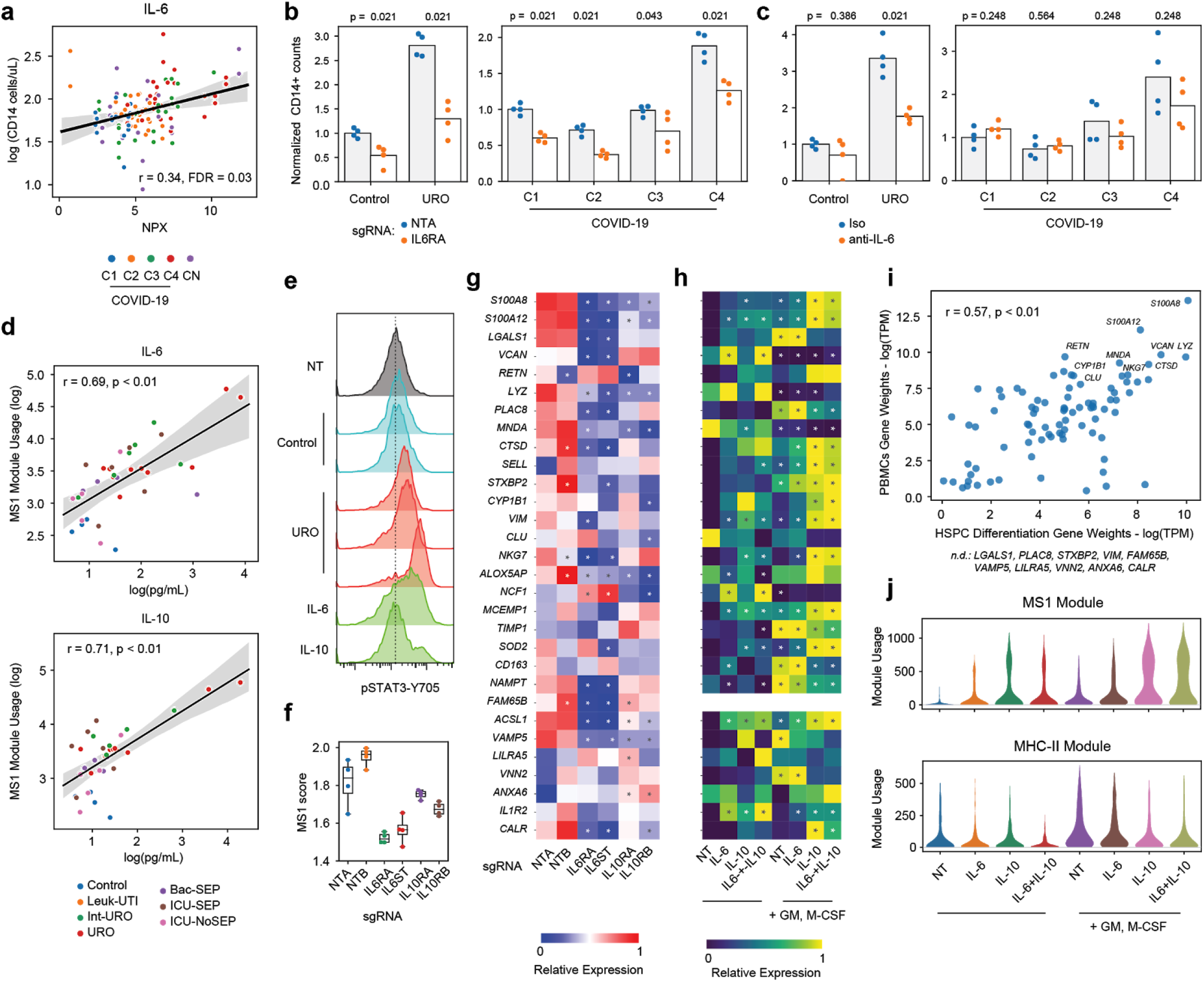
Induction of the MS1 program from CD34+ HSPCs is dependent on IL-6 and IL-10. (a) Correlation between levels of IL-6 in the plasma (NPX, normalized protein expression) and CD34-, CD11b+, CD14+ cell output following incubation of HSPCs with patient plasma (*n =* 61 patients). Line and shadow indicate linear regression fit and 95% confidence interval, respectively. Significance of the correlation (Pearson *r*) was calculated with a two-sided permutation test, corrected for testing of multiple cytokines. (b) CD34-, CD11b+, CD14+ output after incubation of CD34+ HSPCs electroporated with Cas9-RNPs in medium with Control or indicated patient plasma for 7 days. Values are normalized to either the Control (left) or COVID-19 C1 condition for each bone marrow donor. (c) CD34-, CD11b+, CD14+ output after incubation of CD34+ HSPCs in medium containing anti-IL-6 antibody or isotype control with Control or indicated patient plasma for 7 days. Values are normalized to either the Control (left) or COVID-19 C1 condition for each bone marrow donor. Experiments in (b) and (c) were performed on 2 bone marrow donors with 2 technical replicates each using pooled plasma from 5 independent patients or controls for each condition. P-values are calculated using a two-tailed Wilcoxon rank-sum test. (d) Correlation between levels of plasma IL-6 (top) and IL-10 (bottom) and MS1 module usage in each patient (*n* = 40). Line and shadow indicate linear regression fit and 95% confidence interval, respectively. Significance of the correlations (Pearson *r*) were calculated with a two-sided permutation test. (e) Intracellular staining of phosphorylated STAT3 (Y705) in CD34+ HSPCs treated with Control or URO plasma or 100 ng/mL IL-6 or IL-10. Dashed line indicates the median fluorescence for the non-treated control (NT). Results are representative of 2 independent experiments in different bone marrow donors. (f) MS1 scores calculated from bulk RNA-seq of sorted CD14+ cells generated from CD34+ HSPCs electroporated with Cas9-RNPs and treated with sepsis plasma for 7 days. (g) Expression of the top 30 MS1 module genes in sorted CD14+ cells generated from CD34+ HSPCs electroporated with Cas9-RNPs and treated with sepsis plasma for 7 days. Asterisks indicate a significant difference (FDR < 0.1, Wilcoxon rank-sum test, corrected for testing of multiple genes) compared with the non-targeting condition (NTA). In (f,g), *n* = 4 experiments were performed for each condition (2 biological and 2 technical replicates). (h) Expression from scRNA-seq of the top 30 MS1 module genes in CD14-expressing cells generated from CD34+ HSPCs treated with the indicated cytokines (all at 100 ng/mL). Asterisks indicate a significant difference (FDR < 0.1, Wilcoxon rank-sum test, corrected for testing of multiple genes) compared with the non-treated condition (NT). (i) Gene weight correlation between the MS1 modules detected in the cytokine treatment (x-axis) and patient PBMC datasets (y-axis). Significance of the Pearson correlations (r) are calculated with a permutation test. Genes which are not detected (n.d.) in the cytokine treatment module but are among the top 30 for the corresponding module in the PBMC dataset are indicated. (j) Violin plots showing the usage of the MS1 (top) and MHC-II (bottom) modules in CD14-expressing cells across the different cytokine treatment conditions. The experiment in (h-j) was performed on 2 bone marrow donors for each treatment condition; a total of 3,365, 2,986, 2,550, 3,025, 3,194, 3,061, 2,850, and 2,918 cells for each cytokine treatment condition indicated on the plots, respectively, were profiled.

To determine the specific cytokines that induce the differentiation of MS1 cells from HSPCs, we measured the levels of inflammatory cytokines in the plasma of sepsis patients and controls from a previous study^9^. Whereas several cytokines displayed increased concentration in the plasma of sepsis patients (**Supplementary Fig. 6**), we found that both IL-6 and IL-10 levels specifically correlated with the usage of the MS1 module in monocytes (**Figure 3d**), suggesting their specific involvement in its induction, and consistent with the role of both cytokines in MDSC expansion in cancer^38,39^. We found that short-term incubation of HSPCs with URO plasma, but not Control plasma, resulted in phosphorylation of STAT3 (**Figure 3e**), the transcription factor downstream of both cytokines^40,41^. In addition, short term treatment of HSPCs with recombinant IL-6 and/or IL-10 show a dose-dependent up-regulation of the early MS1 genes *S100A8, VCAN*, and *MNDA* (**Supplementary Fig. 7a**) and an increase in S100A8+ progenitor cells (**Supplementary Fig 7b**), similar to the effects observed with sepsis plasma. To further test this dependence, we used CRISPR-Cas9 to knock out the surface receptors for IL-6 (*IL6RA* and *IL6ST*) and IL-10 (*IL10RA* and *IL10RB*) in CD34+ HSPCs before incubation in URO plasma (**Methods, Supplementary Fig. 7c-d**). Knockout of either receptor resulted in the production of CD14+ cells with lower MS1 scores (**Figure 3g**) and reduced expression of several, but not all, MS1 genes (**Figure 3h**). In addition, in these experiments, the CD14+ cells displayed increased surface expression of HLA-DR, consistent with the absence of the MS1 phenotype (**Supplementary Fig. 7e**). Altogether, these results demonstrate that the cytokines IL-6 and IL-10 are necessary for the induction of the MS1 program from HSPCs with sepsis plasma.

To test whether these cytokines are sufficient to induce the MS1 program, we differentiated CD34+ HSPCs with IL-6, IL-10, or both, with and without GM-CSF and M-CSF, which are known growth factors that support the differentiation of HSPCs into monocytes^42^ (**Supplementary Fig 8**). We found that addition of GM-CSF and M-CSF, and to a lesser extent, IL-6 or IL-10, increased the fraction of CD14+ cells produced by HSPCs, but IL-10 decreased the absolute number of cells that are produced (**Supplementary Fig 8g**). Importantly, we found increased expression of several MS1 genes in CD14+ cells generated from HSPCs treated with IL-6 and IL-10 in the presence of GM-CSF and M-CSF (**Figure 3h, Supplementary Fig 8b**). cNMF analysis reveals modules resembling the MS1 and MHC-II modules, both of which correlate significantly with the modules derived from patient PBMC data (Pearson *r* = 0.57 and 0.66, p < 0.01; **Figure 3i, Supplementary Fig 8e-f**). The usage of these modules have opposite trends with cytokine treatment (**Figure 3j**), consistent with their anti-correlation in the PBMC dataset (**Supplementary Fig. 1g**). These results suggest that MS1-like cells can be induced by incubation of HSPCs with the cytokines IL-6 and IL-10 in the presence of GM-CSF and M-CSF, enabling generation of large numbers of these cells for functional studies.

Given their similarity to MDSCs, we hypothesized that MS1 cells suppress the activation of T cells^43^. We generated monocytes from CD34+ HSPCs either with GM-CSF and M-CSF alone (iMono), or with IL-6 and IL-10 (iMS1), and co-incubated the cells with PBMCs activated with anti-CD3 and anti-CD28. We found that incubation with iMS1 cells suppressed the proliferation of T cells, as evidenced by a reduction in the number of cell divisions undergone by CD4 T cells after 4 days (**Figure 4a**). The suppression of T cell proliferation by iMS1 also depended on the ratio of monocytes added to culture (**Figure 4b**). Consistent with this phenomenon, we found that usage of the MS1 module in monocytes correlated negatively with the fraction of CD4+ T cells among total PBMCs in sepsis patients and controls (Pearson *r* = -0.59, p < 0.01; **Figure 4c**). A similar suppressive effect was observed *in vitro* for CD8 T cells, but CD8 T cell levels were not correlated with MS1 module usage (**Supplementary Fig. 9a-c**). These results are consistent with previous studies showing that MDSCs in sepsis patients have immunosuppressive effects on T cells^22,44^.

**Figure 4.**
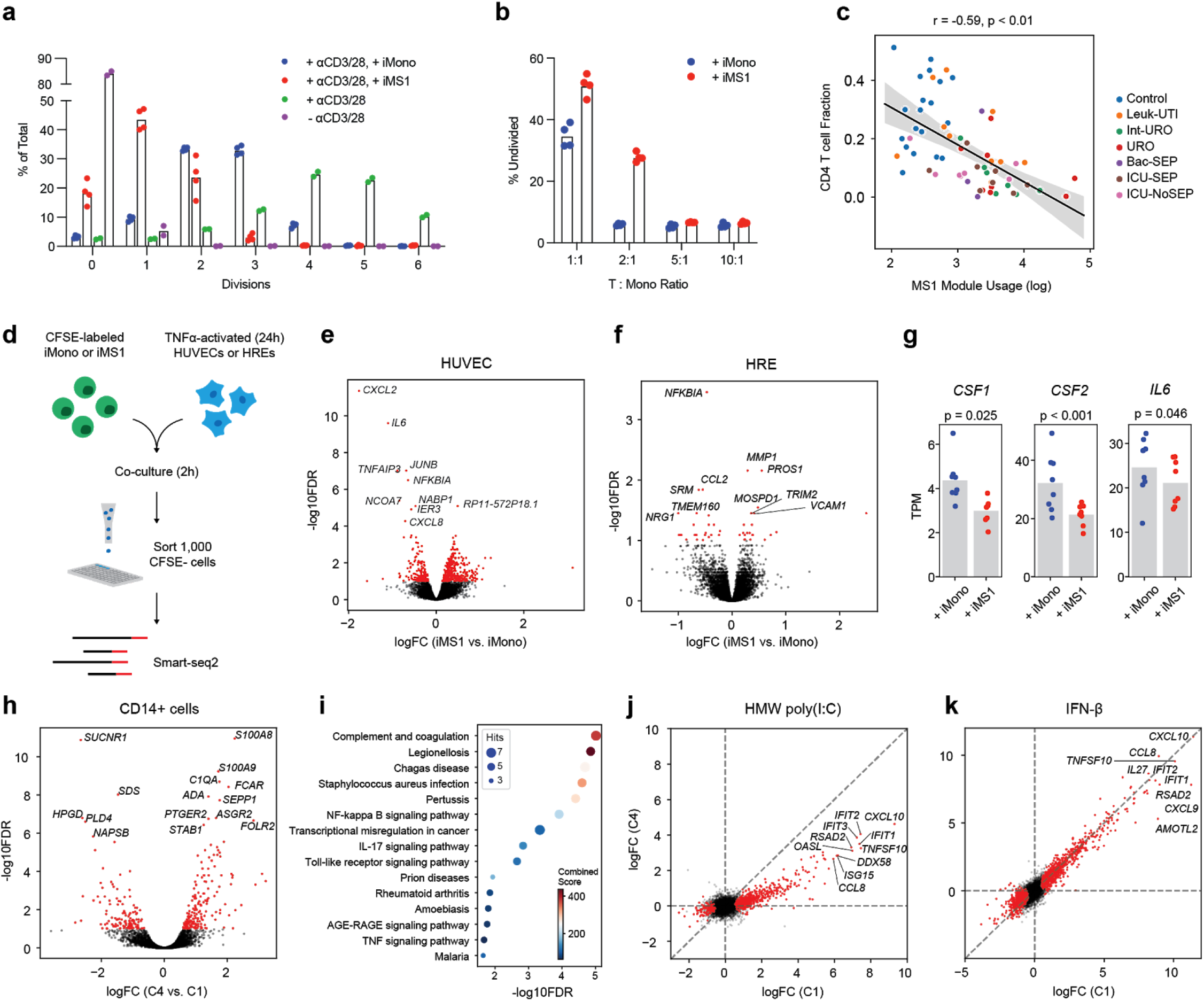
Expression of the MS1 program results in broad immunosuppressive effects. (a) Number of divisions (after 4 days in culture) of CD4 T cells in allogeneic PBMCs activated with CD3 and CD28 antibodies and incubated with either iMono or iMS1 cells generated from CD34+ HSPCs. Percentages are determined by CFSE-dilution (CFSE, carboxyfluorescein succinimidyl ester) and flow cytometry. (b) Fraction of undivided CD4+ T cells in allogeneic PBMCs activated with CD3 and CD28 antibodies and co-incubated with either iMono or iMS1 cells at different ratios. In (a,b), *n* = 4 experiments were performed for each condition (2 bone marrow donors, each with 2 technical replicates). (c) Scatter plot showing the correlation between mean MS1 module usage in monocytes and fraction of CD4+ T cells for each subject (*n* = 65). Line and shadow indicate linear regression fit and 95% confidence interval, respectively. Significance of the correlation (Pearson *r*) was calculated with a two-sided permutation test. (d) Experimental schematic of the co-incubation of iMS1 or iMono with primary human umbilical vein endothelial cells (HUVECs) or human renal epithelial cells (HREs). (e-f) Volcano plot showing differential expression analysis results (exact test) between TNFα-activated HUVECs (e) or HREs (f) co-incubated with either iMono or iMS1 cells generated from CD34+ HSPCs. Genes with FDR < 0.1 are highlighted in red, and the top 10 genes with lowest FDR values are shown. (g) Expression of MS1-inducing cytokines in TNFα-activated HREs co-incubated with either iMono or iMS1 cells generated from CD34+ HSPCs. P values are calculated with an exact test. In (e-g), *n* = 8 experiments were performed for each condition (2 bone marrow donors with 2 biological and 2 technical replicates). (h) Volcano plot showing differential expression analysis results (exact test) between CD14+ cells generated from HSPCs using pooled plasma from C1 or C4 patients. Genes with FDR < 0.1 are highlighted in red, and the top 15 genes with lowest FDR values are shown. (i) Dotplot showing enrichment of pathways (KEGG database) for upregulated genes in CD14+ cells generated with C4 plasma (FDR < 0.1, exact test). (j-k) Scatterplots showing the log2 fold-change of each gene after high molecular weight (HMW) poly-I:C (j) or IFN-β (k) treatment of CD14+ cells generated with pooled plasma from C1 (x-axis) or C4 (y-axis) patients. Genes with FDR < 0.1 in C1 plasma-generated cells are highlighted in red, and the top 10 genes with highest fold-change values are shown. In (h-k), *n* = 4 experiments were performed for each condition (2 bone marrow donors with 2 technical replicates).

Sepsis is a systemic disease; thus, we tested the effects of co-incubating iMS1 cells with cell types from other organs that are commonly dysfunctional in sepsis^45^ (**Methods, Figure 4d**). We found that co-incubation of iMS1 cells with TNFα-activated human umbilical vein endothelial cells (HUVECs) resulted in lower expression of cytokines and chemokines (*CXCL2, CXCL8, IL6*) and down-regulation of the TNFα signalling pathway compared with iMono (**Figure 4e, Supplementary Fig. 9d-e, Supplementary Table 4**). iMS1 cells exerted a similar albeit weaker effect on human renal epithelial cells (HRE), as evidenced by the down-regulation of a number of TNFα signalling pathway genes (*NFKBIA, CCL2, TNIP2, RELB*; **Figure 4f, Supplementary Table 6**). Co-incubation with iMS1 also resulted in decreased expression of *IL6, CSF1*, and *CSF2* in HREs (**Figure 4g**), suggesting a systemic negative feedback loop wherein MS1 suppresses its own induction. These effects were not observed in HUVECs or in HREs incubated in conditioned media from iMS1 or iMono cells, suggesting a cell-contact dependent effect (**Supplementary Fig. 9f**). Of note, co-incubation of HREs with iMS1 cells resulted in the up-regulation of *MMP1*, an important regulator of tissue remodeling and extracellular matrix homeostasis that has been previously shown to be up-regulated during the development of sepsis^46^. Similarly, *PROS1*, the gene encoding protein S, an important regulator of the clotting cascade, was also upregulated in HREs co-incubated with iMS1, suggesting a possible involvement of MS1 cells in sepsis-related coagulopathy^47,48^. These findings demonstrate broad anti-inflammatory effects of MS1 cells and, importantly, highlight other potential functions of monocytes in sepsis beyond their response to pathogens and interaction with other immune cells.

To examine the potential role of MS1 cells in COVID-19, we generated CD14+ cells from HSPCs using plasma from mild (C1) or severe (C4) COVID-19 patients. As expected, the core MS1 gene *S100A8* was the most significantly up-regulated gene in CD14+ cells generated with severe COVID-19 plasma (**Figure 4h, Supplementary Table 5**). Among the positively up-regulated genes, we found enriched pathways related to inflammation and coagulation (**Figure 4i**), both of which contribute to the pathogenesis of severe COVID-19^49,50^. Next, we stimulated C1- and C4-treated populations with high molecular weight (HMW) poly-(I:C), a synthetic RNA analogue, or IFN-β, an important mediator of antiviral responses, and observed weaker induction of various cytokines and interferon-stimulated genes in cells generated with C4 plasma in response to poly-(I:C) (**Figure 4j**), whereas no such effect was observed in response to IFN-β. These results reflect our findings in bacterial sepsis^9^, wherein MS1 cells showed a diminished response to pathogen-associated molecular patterns, and are consistent with a recent report demonstrating functional impairment of myeloid cells in COVID-19 patients^7^. Altogether, these findings suggest that the expansion of MS1 cells in severe SARS-CoV-2 infection is detrimental to the antiviral response and may play a role in the pathogenesis of severe COVID-19.

In this study, we show that expression of the MS1 program is associated with poor outcome in both bacterial and SARS-CoV-2 infections. In addition, we show that systemic cytokines induce emergency myelopoiesis and differentiation of HSPCs into myeloid cells that express the MS1 program, providing a potential explanation for their expansion in bacterial sepsis and severe COVID-19, and a model system through which these cells can be generated and studied in greater detail. Importantly, our study reveals that induction of myelopoiesis and the MS1 program depends on systemic IL-6, a current therapeutic candidate in COVID-19 infection^51^. Our results suggest that the role of IL-6 in infection is complex and more extensive than just the induction of an acute phase response^52^. Given the similarity between MS1 and monocytic MDSCs, we propose that the MS1 gene expression program provides a precise definition for monocytic MDSCs in peripheral blood. In fact, ∼50% genes expressed specifically in monocytic MDSCs vs monocytes, as found in a recent study^5^, overlap with MS1-specific marker genes (**Supplementary Fig. 10**), though our MS1 signature contains many additional genes, suggesting that we have identified a more pure MDSC-like population; however, analysis of additional contexts in which MDSCs are found is warranted^43,54–56^. We also demonstrate that MS1 cells have an anti-inflammatory effect on endothelial and epithelial cells, raising the possibility that MS1 contributes to the pathology of these cell types during bacterial sepsis and severe COVID-19. Whereas these findings suggest several potential roles for MS1 during infections, whether it plays a causal role in the pathogenesis of bacterial sepsis or severe COVID-19 must still be tested. Our study highlights the importance of hematopoietic reprogramming in bacterial sepsis and severe COVID-19 and supports the hypothesis that the interaction of MS1 cells with other cells and tissues impacts pathogenesis of severe infections.

## METHODS

### Bone marrow CD34+ progenitor isolation and culture

Purified CD34+ bone marrow cells from healthy individuals were either purchased directly from StemCell Technologies or isolated from fresh human bone marrow. Bone marrow aspirates anticoagulated with EDTA were purchased from StemExpress and processed within 24 h of isolation. To remove red blood cells (RBC) from the bone marrow, 1X RBC lysis buffer (eBioscience) was added directly at a 10:1 ratio to the sample. After 5 min, the cells were centrifuged at 400g for 5 min and resuspended in 1X RBC lysis buffer to further clear the sample of RBCs. The cells were then centrifuged, resuspended in FACS buffer (1X PBS, 2.5% FBS, 2 mM EDTA, Invitrogen), and purified using human CD34 Microbeads (Miltenyi Biotec). Isolated CD34+ cells were validated using flow cytometry (CD34-BV650, clone 561; BioLegend) to be of >90% purity. Cells were cryopreserved in Cryostor CS10 (Stemcell Technologies) in aliquots of 200,000 cells each. The tubes were kept at −80 °C overnight, then transferred to liquid nitrogen for long-term storage.

For each experiment, CD34+ cells were first thawed and rested for 48 h in SFEM II (StemCell Technologies) with 75 nM StemRegenin 1 (StemCell Technologies), 3.5 nM UM171 (StemCell Technologies), 40 ng/mL SCF, TPO, Flt3L (PeproTech), and 1X penicillin–streptomycin (Gibco). Cells were subsequently cultured in the same medium supplemented with either 20% plasma sterilized through a 0.2 μm filter (Millipore) or 100 ng/mL of the cytokines IL-6, IL-10, GM-CSF, and M-CSF (PeproTech). iMS1 and iMono cells were isolated after 7 d of culture using human CD14 Microbeads (Miltenyi Biotec). Isolated CD14+ cells were validated using flow cytometry (CD14-FITC, clone M5E2; BioLegend) to be of >90% purity.

### Flow cytometry for assessing myeloid output

To assess the number of myeloid cells from plasma incubation experiments, cells were stained with the following panel: CD3-APC (clone HIT3a), CD19-APC (clone HIB19), CD56-APC (clone 5.1H11), CD14-FITC (clone M5E2), CD15-AF700 (clone HI98), CD11b-PE-Cy7 (clone ICRF44), CD34-BV650 (clone 561), and CD38-PE/Cy5 (clone HIT2) (BioLegend). After staining, cells were resuspended in FACS buffer with 2% CountBright beads (Invitrogen) to allow determination of absolute counts during analysis. Flow cytometry data were acquired on a Cytoflex LX (Beckman Coulter) and analyzed using FlowJo v10.1.

### Single-cell RNA sequencing and data analysis

Single cell RNA-seq combined with cell hashing^57^ was performed as previously described^9^. Briefly, cells from multiple culture conditions were labeled with HTO antibodies (BioLegend) and loaded on the Chromium platform using the 3’ v3 profiling chemistry (10X Genomics). Libraries were sequenced to a depth of ∼25,000 reads per cell on a Nextseq 550 (Illumina). The data were aligned to the GRCh38 reference genome using cellranger v3.1 (10X Genomics). Due to their technical incompatibility with droplet-based platforms, scRNA-seq of neutrophils was performed using plate-based Smart-seq2 as previously described^58^.

Single cell data analysis was performed using scanpy^59^ with the same pre-processing and filtering parameters described in a prior publication^9^. To identify the major cell types in the differentiation experiments, we assessed the expression of the following marker genes: HSPCs: *CD34* and *CD38*, monocytes: *CD14* and *LYZ*, neutrophils: *ELANE* and *MPO*, megakaryocytes: *PPBP* and *PF4*. MS1 scores were calculated for the top 30 genes from the MS1 module derived from the sepsis dataset using the ‘score_genes’ in scanpy (ctrl_size = 50, n_bins = 25). RNA velocity analysis was performed using the scVelo package^60^ using the default parameters.

Consensus non-negative matrix factorization analysis was performed as detailed in a previous publication^30^. Briefly, the top 3,000 variable genes for each dataset was first selected to filter the gene expression matrix. NMF was then performed with k=5 to 25 (10 iterations for each k). The number of modules (k) for downstream analysis was selected based on biological interpretability of the modules and stability of the cNMF solution. To ensure that no modules from technical artifacts were analyzed, only gene programs with mean usage >10 across all cells were included for further analysis.

### Bulk data deconvolution and meta-analysis

The gene loading matrix from cNMF analysis of monocytes was used to construct a reference matrix for gene expression deconvolution. To reduce the number of genes in the reference matrix, only the top 1,000 variable genes within the monocyte data were included. Sepsis datasets with survival annotation were obtained from a previously published meta-analysis^33^. Gene expression deconvolution was performed using CIBERSORT^61^ with a no-sum-to-one constraint and absolute scoring. The resulting score matrix was then used as an input to MetaIntegrator^62^, where the effect size of each gene module was visualized using forest plots.

### Sepsis plasma samples

Plasma samples from sepsis patients and controls were obtained from an existing cohort of patients described in a prior publication^9^. Plasma samples were isolated by obtaining the top layer from Ficoll gradient separation of whole blood (diluted 1:1 with 1X PBS) and were centrifuged again at 1000g for 10 min to remove cell debris. Samples were immediately stored at −80 °C.

### COVID-19 plasma samples

COVID-19 plasma samples were obtained from patients entering the Massachusetts General Hospital Emergency department. This study was approved by the Partners Healthcare Institutional Review Board under protocol 2017P001681. Patients presenting to the MGH ED from March through May 2020 with respiratory distress suspected or known to be due to COVID-19 were enrolled. Inclusion criteria were age 18 years or older, clinical concern for COVID-19 upon Emergency Department presentation, and acute respiratory distress with at least one of the following: 1) tachypnea ≥ 22 breaths per minute, 2) oxygen saturation ≤ 92% on room air, 3) a requirement for supplemental oxygen, or 4) positive-pressure ventilation. Patients were categorized based on disease outcomes as follows: C1: non-hospitalized, C2: hospitalized without intensive care, C3: hospitalized and admitted to intensive care, C4: hospitalized and eventually died, and CN: SARS-CoV-2 negative patients. Blood samples were collected upon hospital presentation. Plasma was obtained as described above, but using whole blood diluted 1:2 in RPMI-1640. Prior to use in experiments, plasma samples were quickly thawed at 37°C and incubated for 1 h at 53°C to inactivate viral particles. For the knockout and scRNA-seq experiments, plasma samples were pooled across each patient category.

### Blood samples for neutrophil sorting and scRNA-seq

Samples for neutrophil scRNA-seq were from patients enrolled in the Brigham and Women’s Hospital’s (BWH); the criteria for patient recruitment for this cohort are described elsewhere^63,64^. Control samples consisted of blood samples from age, gender, and ethnicity-matched healthy controls obtained from Research Blood Components (MA, USA).

To maintain viability of neutrophils for scRNA-seq, fresh blood was collected in EDTA Vacutainer tubes (BD Biosciences) and processed within 4 h. To remove red blood cells (RBC) from the sample, 1X RBC lysis buffer (eBioscience) was added directly at a 10:1 ratio to the sample. After 5 min, the cells were centrifuged at 400g for 5 min and resuspended in 1X RBC lysis buffer to further clear the sample of RBCs. The remaining cells were stained with a general panel: DAPI, CD3-APC (clone HIT3a), CD19-APC (clone HIB19), CD56-APC (clone 5.1H11), CD14-FITC (clone M5E2), CD15-AF700 (clone HI98), CD11b-PE-Cy7 (clone ICRF44) (BioLegend). Single neutrophils were sorted using an SH800 cell sorter (Sony) into 10 μL TCL buffer (Qiagen) with 1% β-mercaptoethanol (BME, Sigma) in individual wells of a 96-well plate.

### Bulk RNA-seq processing and data analysis

Bulk RNA-seq was performed using Smart-Seq2^65^ with minor modifications, as described previously^66^, using 1,000 cells as input. All RNA-seq libraries were sequenced with 38 × 38 paired-end reads using a NextSeq (Illumina). RNA-seq libraries were sequenced to a depth of >2 million reads per sample. STAR was used to align sequencing reads to the UCSC hg19 transcriptome and RSEM was used to generate an expression matrix for all samples. Both raw count and transcripts per million data were analyzed using edgeR and custom python scripts.

### Intracellular protein staining

Intracellular staining with S100A8-PE (clone REA917, Miltenyi Biotec) was performed using the Cytofix/Cytoperm kit (BD Biosciences) following the manufacturer’s protocol. For staining of phosphorylated STAT3 (Y705), cells were first fixed with 4% paraformaldehyde (Electron Microscopy Sciences) for 15 min at room temperature. The cells were then washed twice with 1X PBS and resuspended in 95% ice-cold methanol and left at -20°C overnight. The permeabilized cells were then stained with a pSTAT3-Y705 antibody (clone 13A3-1, BioLegend) for 30 mins on ice. Flow cytometry data were acquired on a Cytoflex LX (Beckman Coulter) and analyzed using FlowJo v10.1.

### CRISPR-Cas9 editing of CD34+ HSPCs

Cas9 protein, pre-designed guide RNAs targeting *IL6ST, IL6R, IL10RA*, and *IL10RB* and non-targeting guide RNAs (from GeCKO v2 library) were purchased from Integrated DNA Technologies. RNP complexes were assembled by combining 2.1 μL 1X PBS, 1.2 μL 100 μM gRNA, and 1.7 μL 10 μg/mL Cas9 protein and incubating at room temperature for 15 min. The complexes were added to 50,000 - 100,000 CD34+ HSPCs resuspended in 20 μL P3 (Lonza) and electroporated (program code DZ-100) using the 4D-Nucleofector system (Lonza). After electroporation, the cells were immediately transferred to 500 μL of HSPC media and rested for 48 h. Knockout efficiency in CD34+ HSPCs was assessed after 48 h via flow cytometry using the following panel: CD34-BV650 (clone 561), CD38-PE/Cy5 (clone HIT2), CD126-APC (clone UV4), CD210-PECy7 (clone 3F9) (BioLegend).

### Plasma protein measurements

Samples from sepsis patients and controls were thawed and analyzed in parallel using the Legendplex Human Inflammation for IFN-α2, IFN-γ, IL-1β, IL-6, IL-10, TNFα and IL-18, and Human Hematopoietic Stem Cell Panels for M-CSF and GM-CSF (BioLegend). Flow cytometry data were acquired on a Cytoflex LX (Beckman Coulter) and analyzed using FlowJo v10.1.

Samples from COVID-19 patients and controls were analyzed using a commercially available multiplexed proximity extension assay (Olink Proteomics).

### T cell co-culture and proliferation assay

T cell co-culture was performed as previously described^67^ with minor modifications. Briefly, tissue-culture plates were coated with 5 μg/mL purified anti-CD3 (clone HIT3a, Biolegend) at 4°C overnight and subsequently washed twice with 1X PBS. PBMCs from a healthy donor (Research Blood Components) were labelled with CFSE (Invitrogen) following the manufacturer’s protocol. PBMCs were resuspended in SFEM II (StemCell Technologies) with 5 μg/mL purified anti-CD28 (clone CD28.2, BioLegend) and plated at a density of 1M cells/mL. Isolated iMS1 or iMono cells were added at different ratios as indicated. The cells were left in culture for 3-4 days, with media replenished after 2 days. At the end of incubation, the cells were stained with CD3-AF700 (clone OKT3), CD4-APC (clone OKT4), and CD8a-PE (clone RPA-T8) (BioLegend) to determine the amount of CFSE dilution within the T cells.

### HUVEC and HRE co-culture and flow sorting

Primary HUVECs and HREs were purchased from Lonza and cultured in EGM-2 and REGM, respectively. To improve cell viability, HUVECs were cultured in tissue culture plates pre-coated with Matrigel (Corning) diluted 1:100 in EBM-2 (Lonza). Cells were used for experiments within 3-5 passages. Prior to co-culture, HUVECs and HREs were treated with 10 ng/mL TNFα (PeproTech) for 24 h. For the co-culture experiments, CFSE-labelled iMono or iMS1 cells were added at a 1:1 ratio to a confluent monolayer of HUVECs or HREs and incubated for 2 h. The cells were washed 3x with 1X PBS and detached by adding 1X Accutase (Innovative Cell Technologies). The cells were transferred to FACS buffer after 15 mins, and 1,000 CFSE-negative cells were sorted into 10 μL TCL buffer (Qiagen) with 1% BME (Sigma) for bulk RNA-seq. Conditioned media was prepared by incubating iMS1 or iMono cells at 0.5 M cells/mL in EGM-2 or REGM for 24 h overnight.

### Stimulation of CD14+ cells generated from COVID-19 plasma

C1 and C4-plasma generated cells were isolated after 7 d of culture using human CD14 Microbeads (Miltenyi Biotec). Isolated CD14+ cells were validated using flow cytometry (CD14-FITC, clone M5E2; BioLegend) to be of >90% purity. Cells were stimulated for 4h with 100 ng/mL IFN-β (PeproTech) or 1 μg/mL HMW poly-I:C (Invivogen). After stimulation, cells were washed twice with FACS buffer and resuspended in 20 μL TCL buffer (Qiagen) with 1% BME (Sigma) for bulk RNA-seq.

## ACKNOWLEDGEMENTS

We thank Matteo Gentili, Bingxu Liu, Thomas Eisenhaure, Anna Le, Mohammad Najia, Rebecca Carlson, Joshua Peters, Joshua Elacqua, and other members of the Blainey and Hacohen Labs (Broad Institute) for helpful discussions. We thank Alec Schmaier and Samir Parikh (Beth Israel Deaconess Medical Center) for advice on endothelial cell experiments. We also thank the Broad Flow Cytometry core for assistance in cell sorting experiments and the Broad Genomics Platform for assistance in sequencing. We are grateful to Drs. Laura Fredenburgh, Paul Dieffenbach, and Sam Ash for assistance with patient phenotyping at Brigham and Women’s Hospital.

This work was supported by NIH NIAID U24 AI118668 (N.H. and P.C.B.). N.H. was supported by the David P. Ryan, MD Endowed Chair in Cancer Research, R.P.B. was supported by a Mentored Clinical Scientist Research Career Development Award from the NIH (1K08AI119157-04), M.B.G., M.R.F., and R.P.B. were supported by a grant from the Executive Committee on Research, Massachusetts General Hospital, M.B.G. was supported by a grant from the American Lung Association. A.S. was supported by a T32 institutional training grant from NIH (T32 GM007592).

## AUTHOR CONTRIBUTIONS

M.R. designed and performed experiments, and analyzed the data. K.B. performed neutrophil sorting and measurement of cytokines in plasma samples. A.S. analyzed the neutrophil data. A.M. analyzed plasma from COVID-19 samples. M.R.F., R.P.B., M.B.G. designed the MGH bacterial sepsis and COVID-19 clinical cohorts and supervised patient enrollment, specimen collection, and performed clinical adjudications. K.R.K., A.C.V., M.S.-F. supervised specimen collection for the MGH COVID-19 clinical cohort. L.A.C., D.T.H., B.D.L., R.M.B. supervised patient enrollment and specimen collection at BWH. M.P.V., M.E.B performed specimen collection at BWH. M.R., P.C.B., and N.H. conceived the study. N.H., and P.C.B. supervised the study. M.R., P.C.B, and N.H. prepared the manuscript; all authors reviewed and edited the final manuscript.

## DATA AND CODE AVAILABILITY

Gene expression data will be made available upon publication.

## SUPPLEMENTARY FIGURES

**Supplementary Figure 1.**
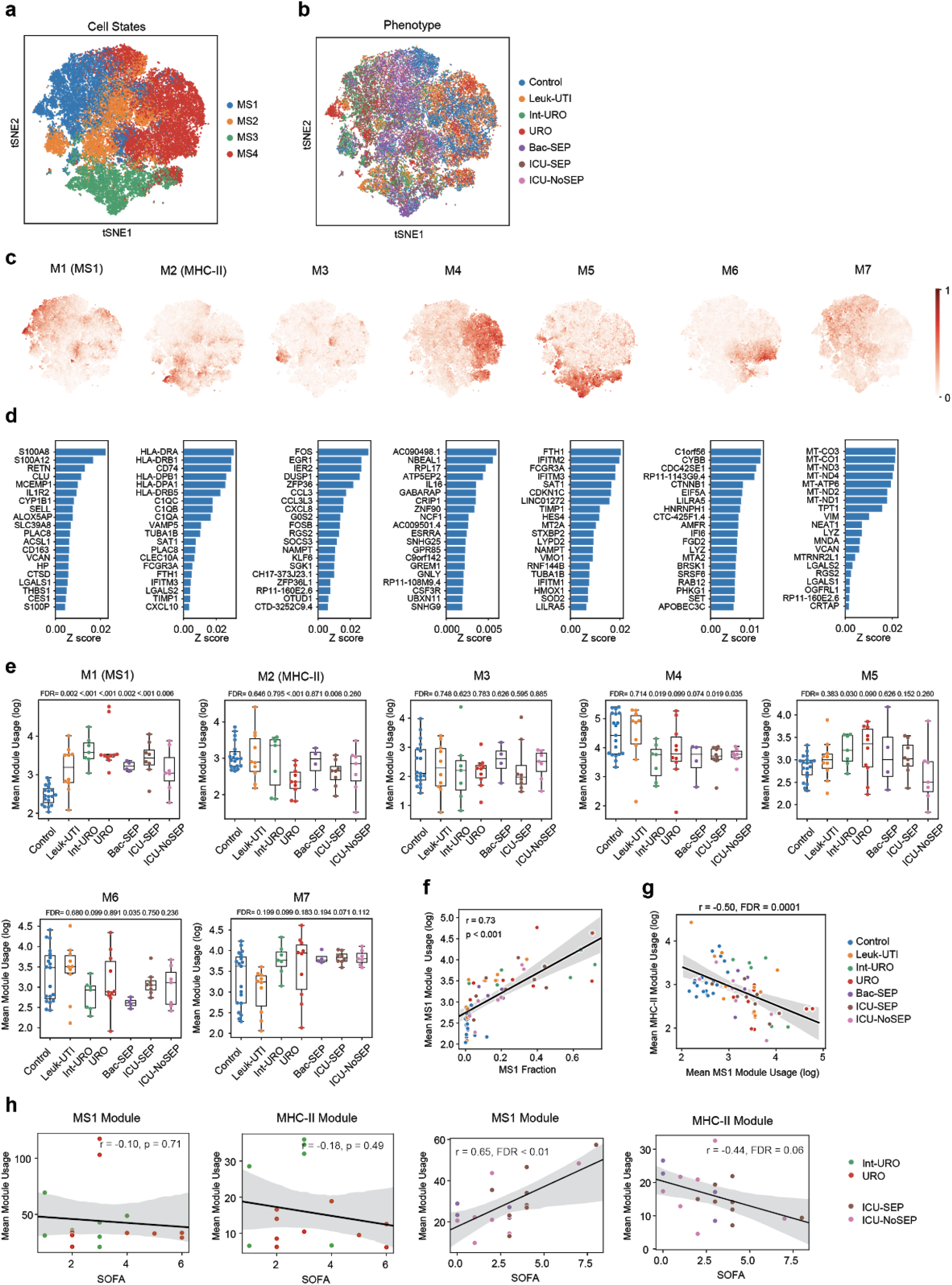
Characterization of gene expression modules in sepsis monocytes. (a-b) tSNE plot of monocytes from sepsis patients and controls colored by (a) cell state and (b) cohort. (c) Relative usage of gene expression module unbiasedly derived from the monocyte scRNA-seq data. (d) Genes with the top 10 loading in each gene expression module. (e) Mean usage of each gene module in monocytes for each patient type. FDR values are shown when comparing each disease state with healthy controls (two-tailed Wilcoxon rank-sum test, corrected for testing of multiple modules). Boxes show the median and IQR for each patient cohort, with whiskers extending to 1.5 IQR in either direction from the top or bottom quartile. (f) Scatterplot showing correlation between mean MS1 gene module usage in monocytes and MS1 fractions for each patient. Line and shadow indicate linear regression fit and 95% confidence interval, respectively. Significance of the correlations (Pearson r) were calculated with a two-sided permutation test. (g) Scatterplot showing correlation between mean MS1 and MHC-II gene module usage in monocytes for each patient. Line and shadow indicate linear regression fit and 95% confidence interval, respectively. Significance of the correlations (Pearson r) were calculated with a two-sided permutation test and corrected for multiple comparison of modules. (h) Scatter plots showing the correlation between mean module usage in monocytes and SOFA scores for each subject. Line and shadow indicate linear regression fit and 95% confidence interval, respectively. Significance of the correlations (Pearson *r*) were calculated with a two-sided permutation test. Detailed description of the patient cohorts and number of cells and patients are outlined in Reyes, et al^9^. Control, healthy controls; Leuk-UTI, urinary tract infection with leukocytosis; Int-URO, intermediate urosepsis; URO, urosepsis; Bac-SEP, sepsis with confirmed bacteremia; ICU-SEP, intensive care with sepsis; ICU-NoSEP, intensive care without sepsis; SOFA, sequential organ failure assessment.

**Supplementary Figure 2.**
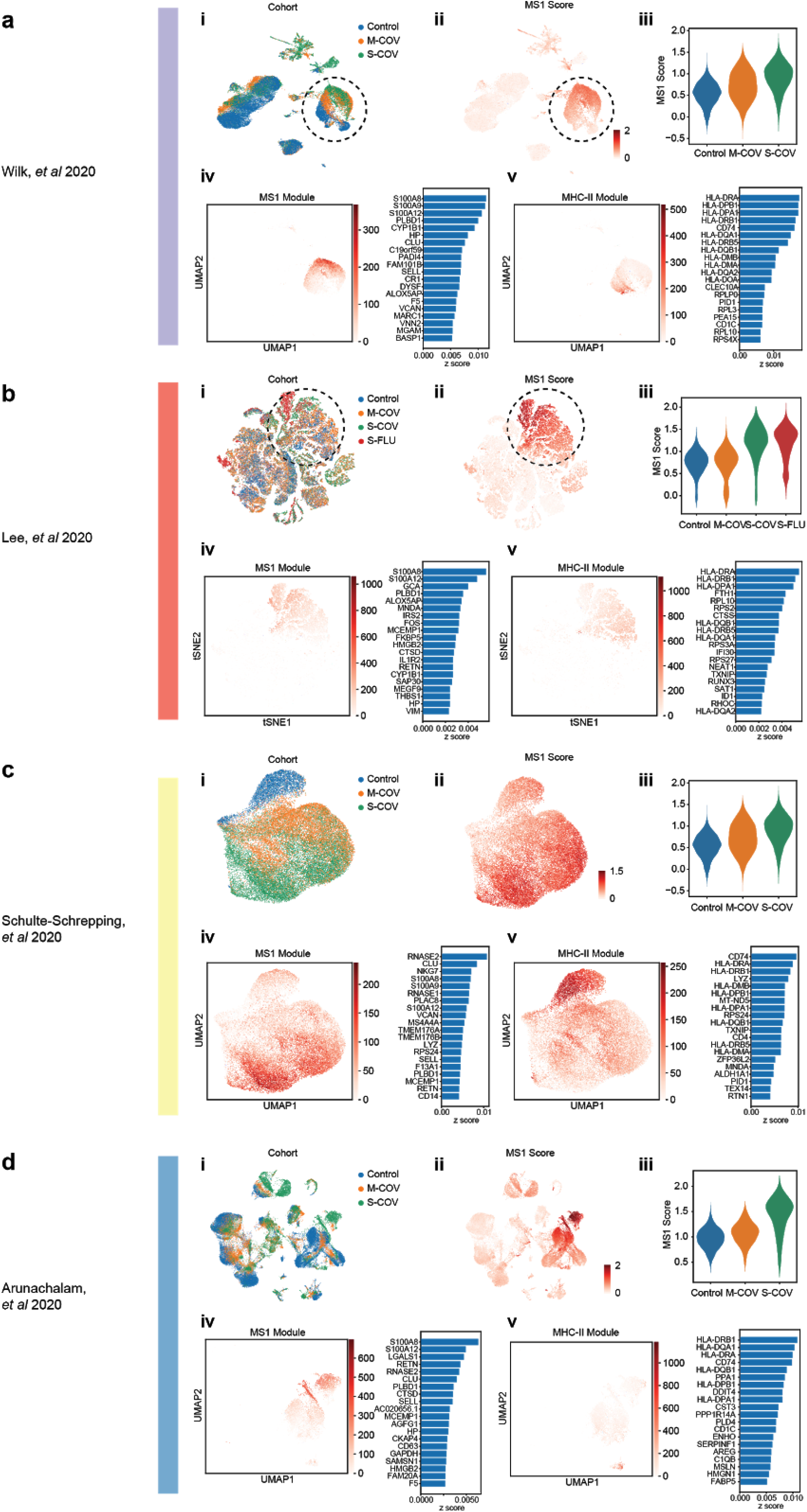
Analysis of MS1 and MHC-II gene expression modules in COVID-19 blood scRNA-seq datasets. Independent analysis of four datasets: (a) Wilk, *et al* 2020, (b) Lee, *et al* 2020, (c) Schulte-Schrepping, *et al* 2020, and (d) Arunachalam, *et al* 2020. For each dataset: (i-ii) UMAP or tSNE projection of the cells colored by (i) patient condition and (ii) MS1 scores. (iii) Violin plot of MS1 scores for CD14-expressing cells from each condition. (iv-v) Left, UMAP or tSNE plot, showing usage of the MS1 and MHC-II gene expression modules unbiasedly derived from each dataset. Right, Barplots of the top 20 genes with the highest z-score loading in each module. Detailed description of the patient cohorts and number of cells and patients for each dataset in (d,e) are outlined in the corresponding publications^7,28,34,35^. M-COV, mild COVID-19; S-COV, severe COVID-19, S-FLU, severe influenza A.

**Supplementary Figure 3.**
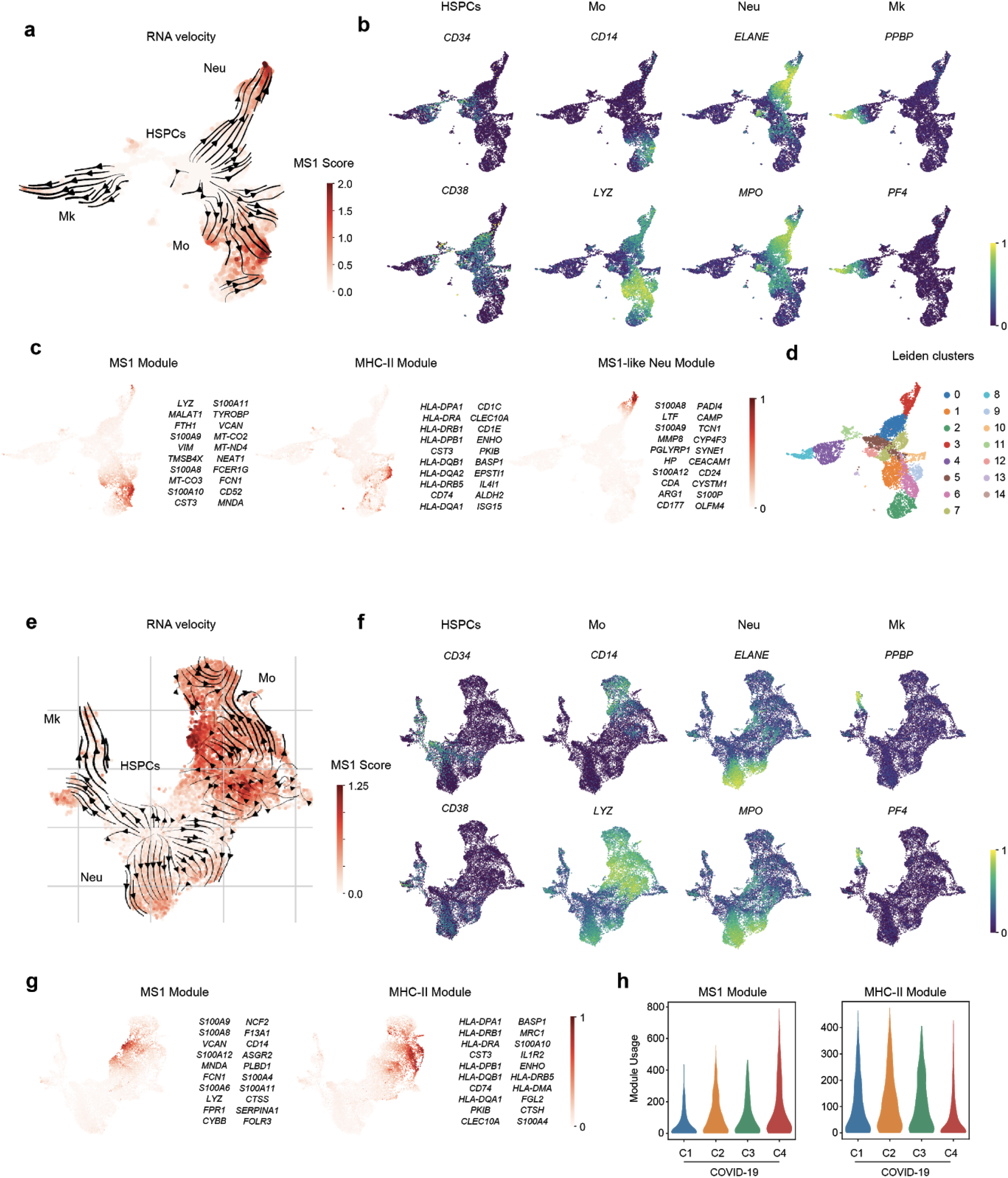
scRNA-seq of plasma-treated CD34+ HSPCs. (a-g) UMAP projection of differentiated cells from CD34+ HSPCs incubated with (a-d) sepsis or (e-g) COVID-19 plasma and corresponding controls. Cells are colored by (a,e) MS1 scores, (b,f) relative expression of HSPC, monocyte (Mo), neutrophil (Neu), and megakaryocyte (Mk) marker genes, (c,g) relative usage of the MS1 and MHC-II modules, and (d) Leiden clusters. Stream plots showing inferred trajectories from RNA velocity are overlaid in (a,e). Top 20 genes in each module are listed in (c,g). (h) Violin plots showing the usage of the MS1 (left) and MHC-II (right) modules in CD14-expressing cells across the different plasma treatment conditions. The experiment in (a-d) was performed on 2 healthy bone marrow donors with 2 plasma donors for each condition; a total of 3,039 and 5,254 cells were profiled for Control and URO plasma treatment, respectively. The experiment in (e-h) was performed on 2 healthy bone marrow donors with pooled plasma from all donors in (c); a total of 4,449, 4,591, 3,129 and 3,711 cells were profiled for C1-C4 plasma, respectively.

**Supplementary Figure 4.**
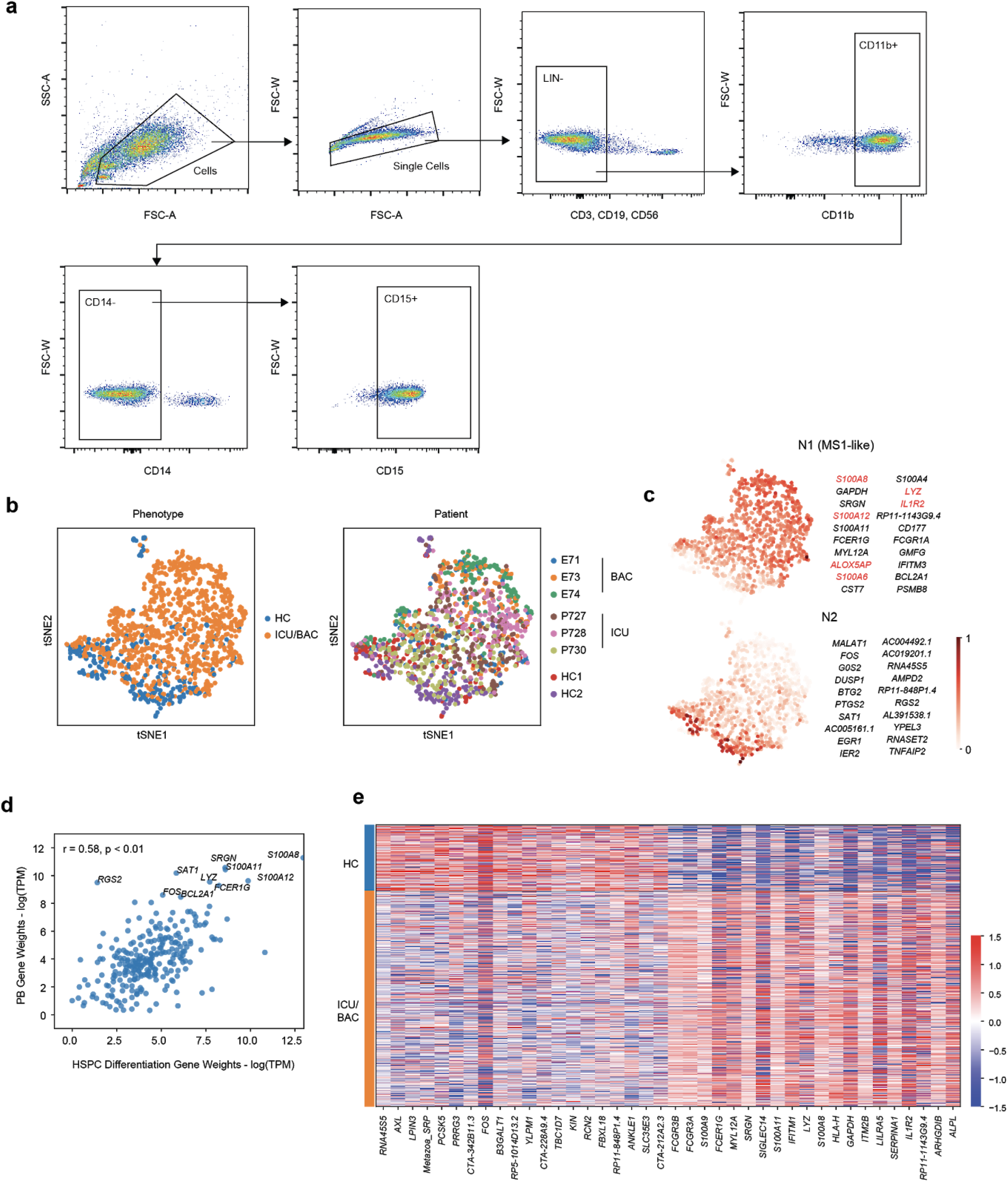
scRNA-seq of neutrophils sorted from critically-ill and bacteremic patients. (a) Gating strategy for single cell sorting of neutrophils. (b) tSNE plot (*n* = 2,168 cells) of neutrophil scRNA-seq data colored by cohort (left) and individual patients (right). (c) Relative usage of two gene modules unbiasedly derived from the neutrophil data (by cNMF). The top 20 genes in each module are listed. Genes in module N1 that are also among the top 30 genes in the MS1 module are highlighted in red. (d) Gene weight correlation between the MS1-like modules detected in neutrophils from the plasma-incubation (x-axis) and patient datasets (y-axis). Significance of the correlations (Pearson *r*) are calculated with a permutation test. (e) Heatmap showing the top 20 differentially expressed genes (Wilcoxon rank-sum test, FDR < 0.01) between neutrophils sorted from critically-ill patients and healthy controls. BAC, bacteraemic patient; ICU, patient in intensive care.

**Supplementary Figure 5.**
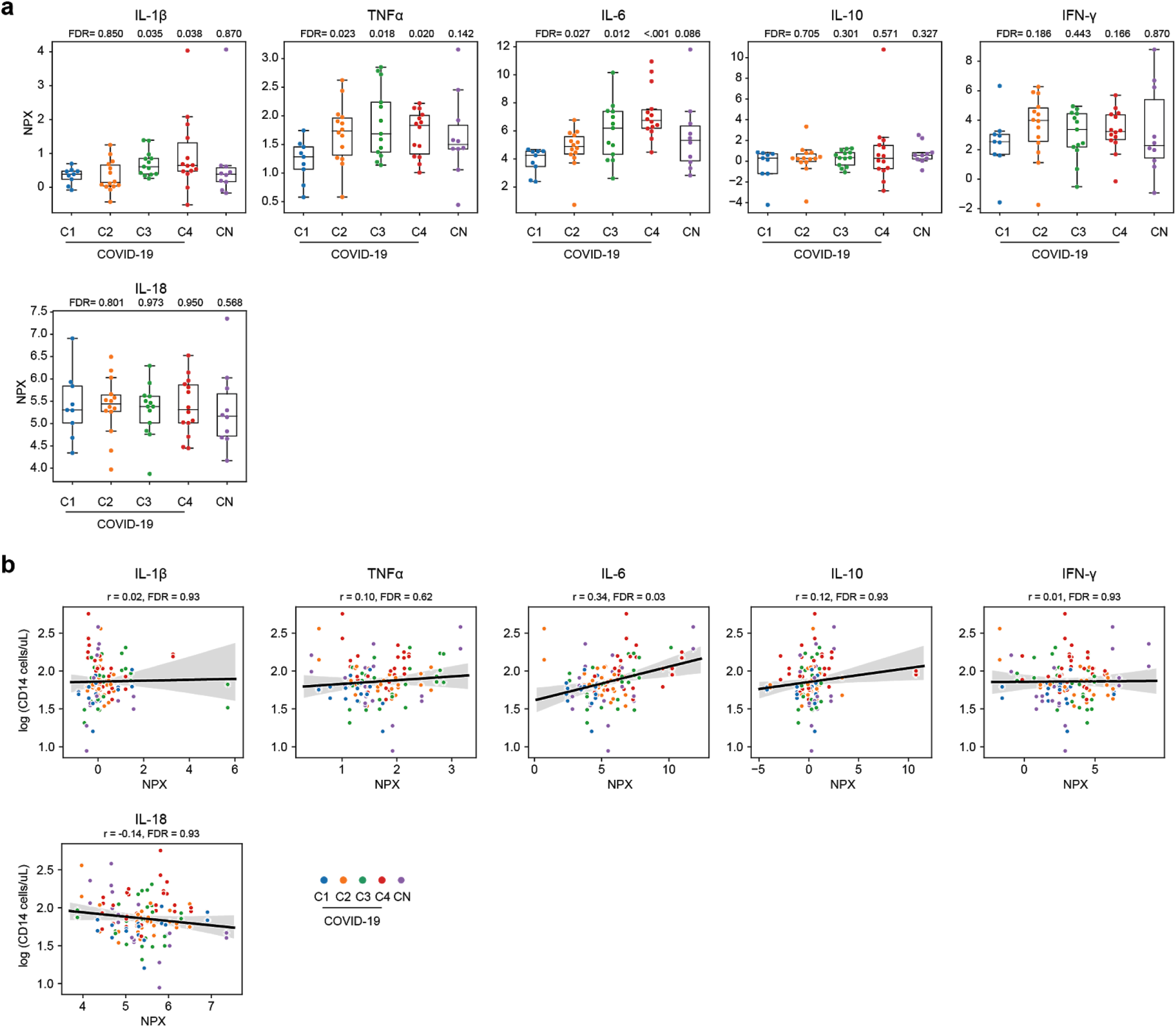
Plasma inflammatory cytokine levels in COVID-19 patients and correlation with cellular output. (a) Boxplots showing NPX (normalized protein expression) levels of inflammatory cytokines in plasma for each patient (*n* = 61). FDR values are shown when comparing each disease state with healthy controls (two-tailed Wilcoxon rank-sum test, corrected for testing of multiple cytokines). Boxes show the median and IQR for each patient cohort, with whiskers extending to 1.5 IQR in either direction from the top or bottom quartile. (b) Correlation between cytokine levels and CD14 cell output after CD34+ HSPCs are incubated with plasma for 7 days. Line and shadow indicate linear regression fit and 95% confidence interval, respectively. Significance of the correlations (Pearson r) were calculated with a two-sided permutation test.

**Supplementary Figure 6.**
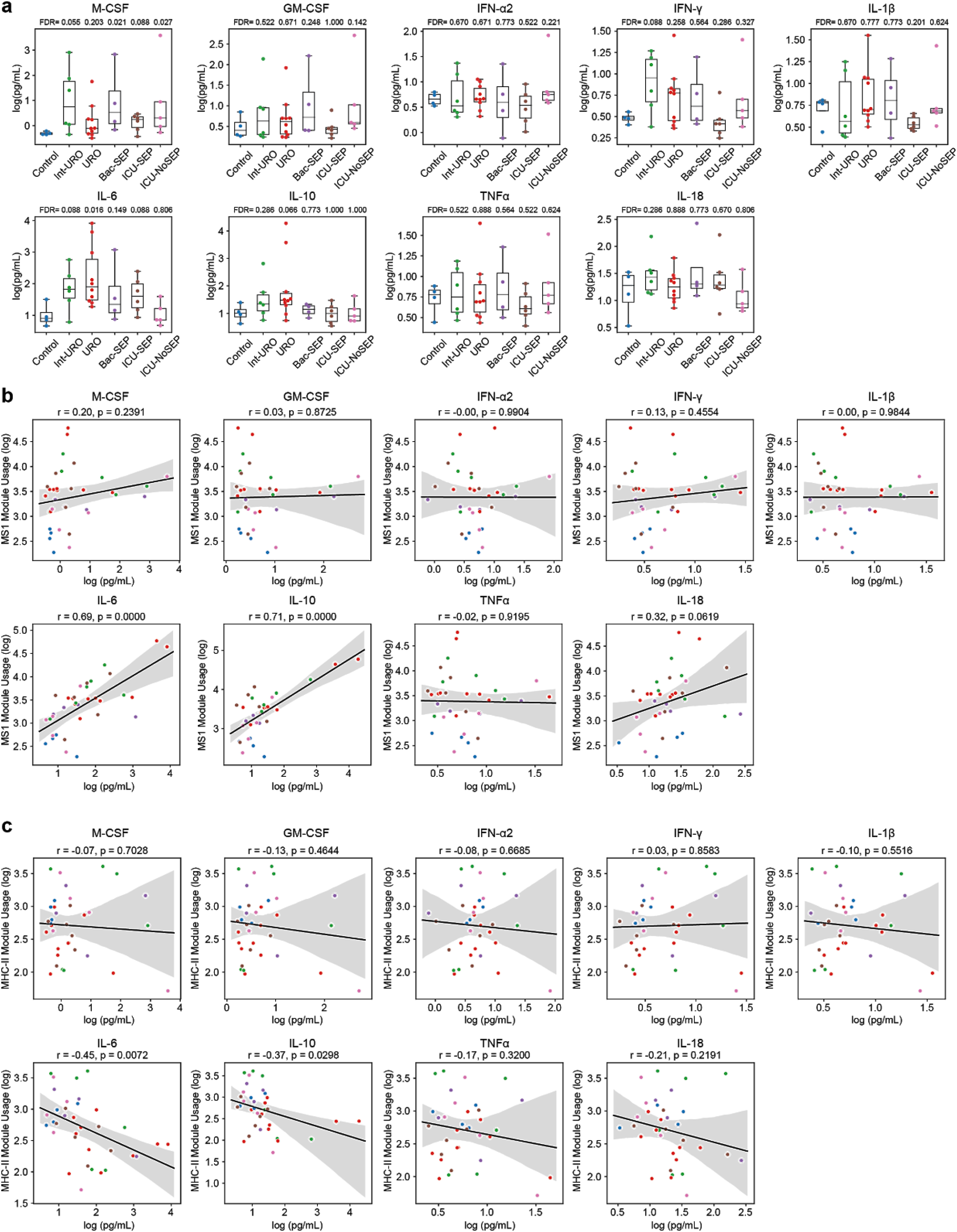
Plasma inflammatory cytokine levels in sepsis patients and correlations with monocyte gene expression modules. (a) Boxplots showing levels of indicated inflammatory cytokines in plasma for each patient (*n* = 40 total). FDR values are shown when comparing each disease state with healthy controls (two-tailed Wilcoxon rank-sum test, corrected for testing of multiple cytokines). Boxes show the median and IQR for each patient cohort, with whiskers extending to 1.5 IQR in either direction from the top or bottom quartile. (b-c) Correlations between level of each indicated cytokine and MS1 (b) or MHC-II (c) module usage in each patient (*n* = 40 total). Line and shadow indicate linear regression fit and 95% confidence interval, respectively. Significance of the correlations (Pearson r) were calculated with a two-sided permutation test.

**Supplementary Figure 7.**
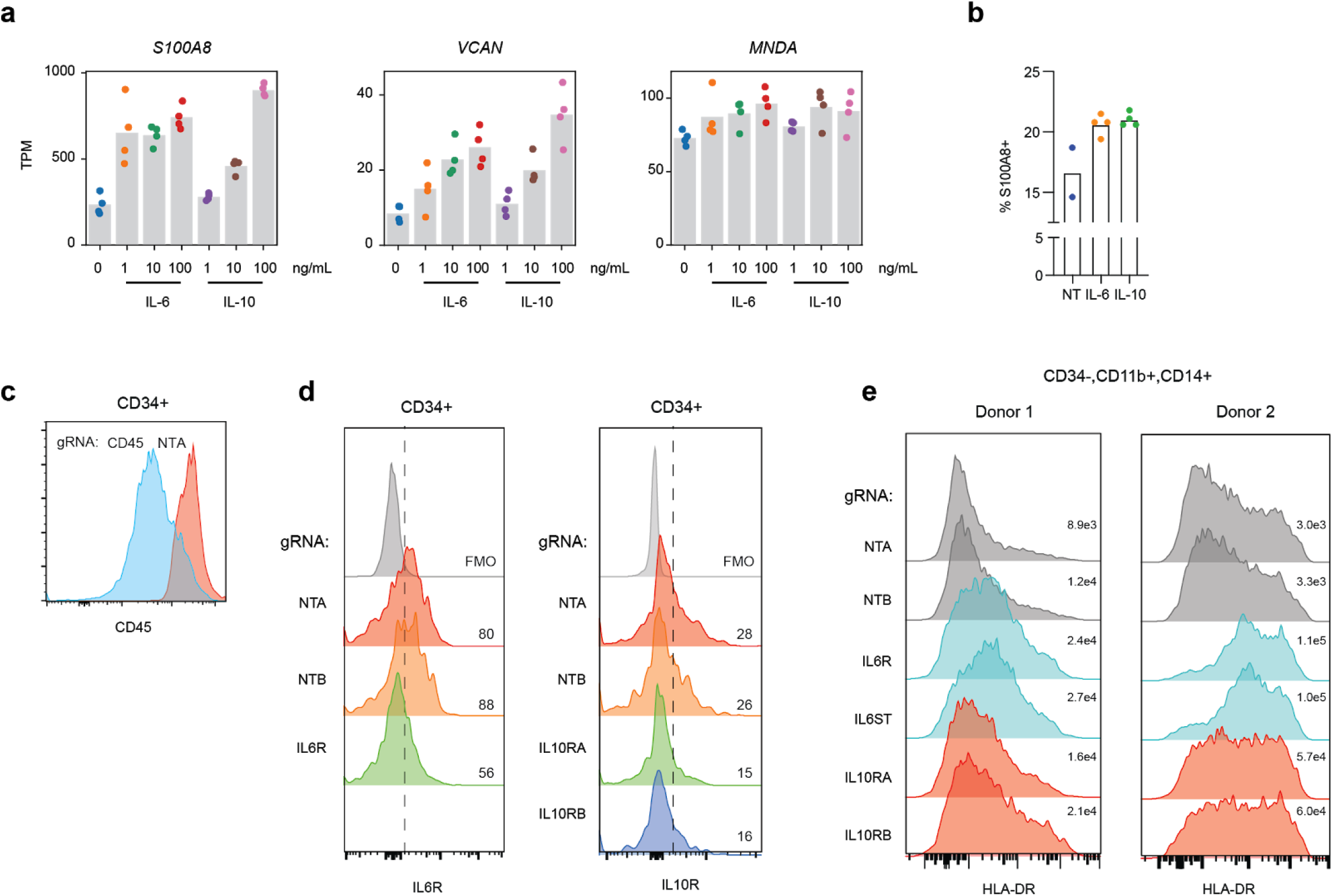
Short term stimulation and Cas9 RNP-based knockouts of CD34+ HSPCs. (a) Expression of ‘early’ MS1 genes in CD34+ HSPCs after treatment with IL-6 or IL-10 for 24 h. Experiments were performed with *n* = 4 for each condition (2 bone marrow donors with 2 technical replicates). (b) S100A8 intracellular staining of CD34+ HSPCs treated with 100 ng/mL IL-6 or IL-10 for 24 h. (c) CD45 surface expression levels in CD34+ HSPCs electroporated with either CD45 or NTA RNPs. (d) IL6R (left) and IL10R (right) expression in CD34+ HSPCs electroporated with the indicated gRNA-RNPs. Numbers indicate the percentage of cells that are IL6R+ or IL10R+, determined by gating on the full-minus-one (FMO) control. Results are representative of 2 independent experiments in different bone marrow donors. (e) Expression of HLA-DR in CD14+ cells after differentiation of RNP-electroporation HSPCs with 20% sepsis plasma for 2 bone marrow donors. Median fluorescence intensities are indicated for each sample.

**Supplementary Figure 8.**
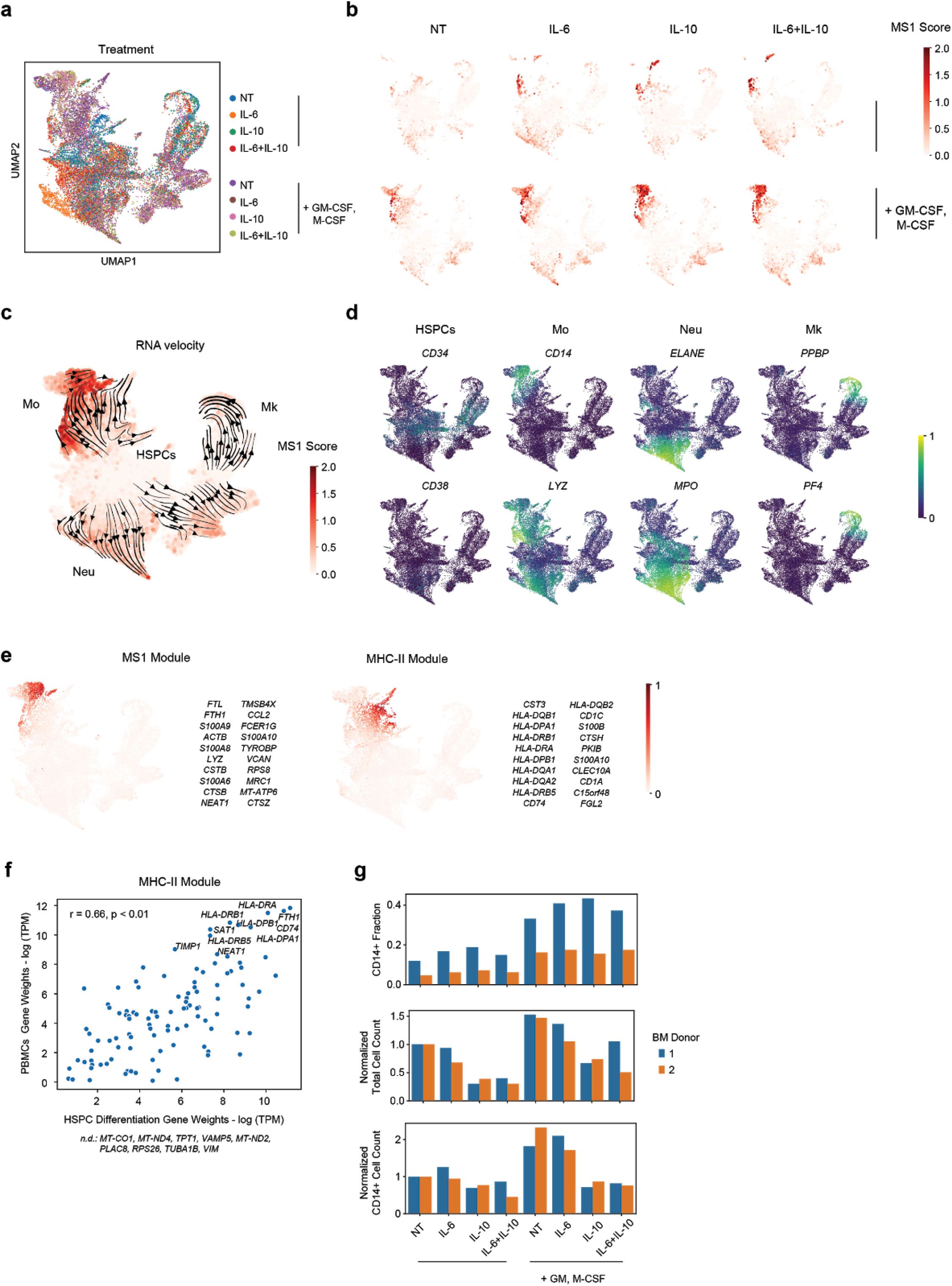
scRNA-seq of cytokine-differentiated HSPCs. (a) UMAP projection of differentiated cells from CD34+ HSPCs incubated with cytokines (all at 100 ng/mL). Cells are colored by cytokine conditions. (b) Individual UMAP projections of each cytokine condition with cells colored by MS1 scores. (c) Combined UMAP projection of all cytokine conditions with cells colored by MS1 scores. Stream plots showing inferred trajectories from RNA velocity are overlaid. Mo, monocytes; Neu, neutrophils; Mk, megakaryocytes. (d) Relative expression of HSPC, monocyte, neutrophil, and megakaryocyte marker genes in (a). (e) Relative expression of the MS1 and MHC-II modules. The top 20 genes in each module are listed. (f) Gene weight correlation between the MHC-II modules detected in the cytokine treatment (x-axis) and patient PBMC datasets (y-axis). Significance of the Pearson correlation (r) is calculated with a permutation test. Genes which are not detected (n.d.) in the cytokine treatment module but are among the top 30 for the corresponding module in the PBMC dataset are indicated. (g) Fraction of CD14+ cells (top), total number of cells (middle), and total number of CD14+ cells (bottom) after incubation of HSPCs in the indication cytokines. Cell counts were normalized to the NT condition (without GM/M-CSF) for each bone marrow donor. The experiment was performed on 2 bone marrow donors for each treatment condition; a total of 3,365, 2,986, 2,550, 3,025, 3,194, 3,061, 2,850, and 2,918 cells for each cytokine treatment condition indicated on the plots, respectively, were profiled.

**Supplementary Figure 9.**
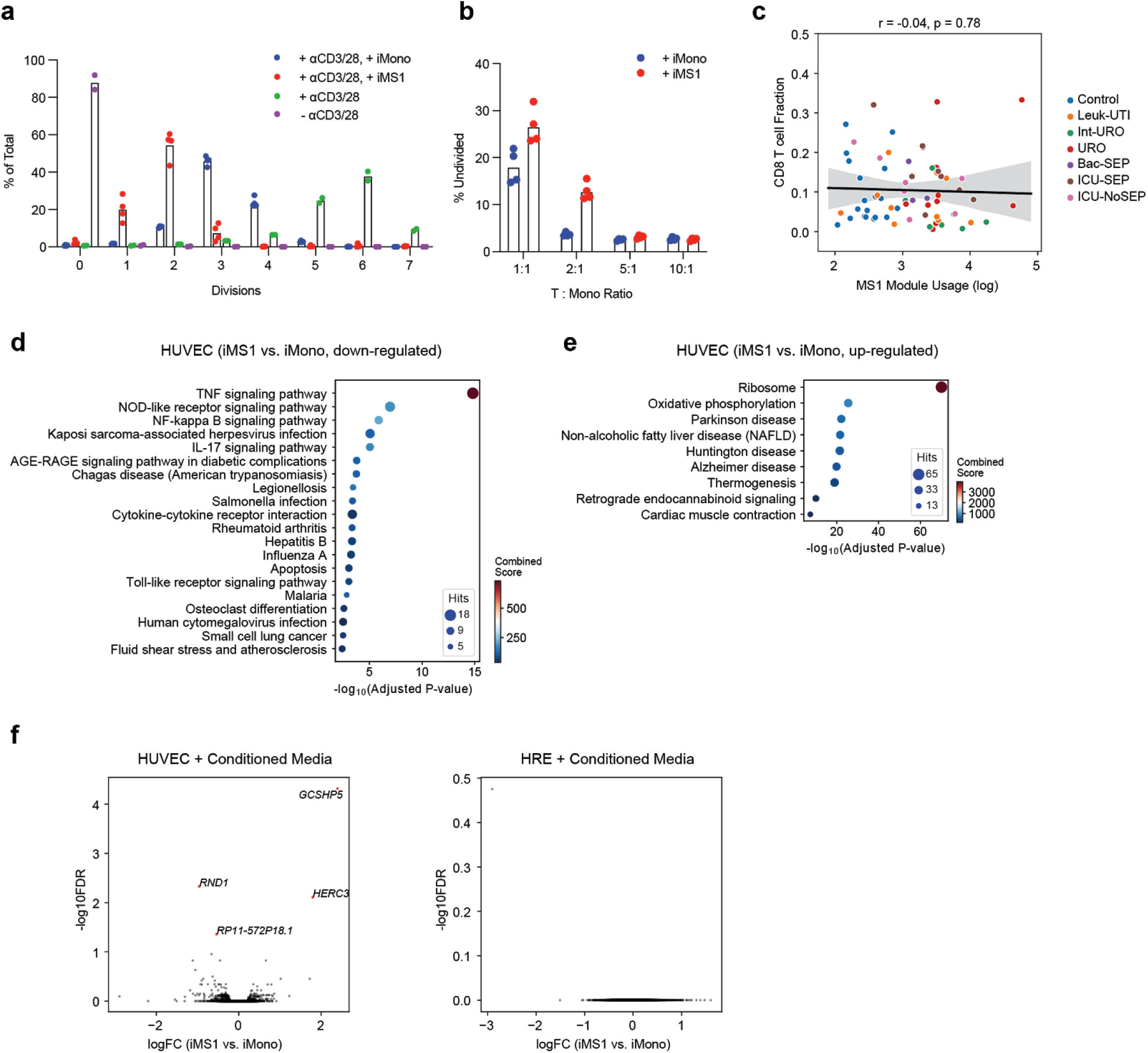
iMS1 co-incubation experiments. (a) Number of divisions (after 4 days in culture) of CD8 T cells in allogeneic PBMCs activated with CD3 and CD28 antibodies and incubated with either iMono or iMS1 cells generated from CD34+ HSPCs. Percentages are determined by CFSE-dilution and flow cytometry. (b) Fraction of undivided CD8 T cells in allogeneic PBMCs activated with CD3 and CD28 antibodies and co-incubated with either iMono or iMS1 cells at different ratios. In (a,b), *n* = 4 experiments were performed for each condition (2 bone marrow donors with 2 technical replicates). (c) Scatter plot showing the correlation between mean MS1 module usage in monocytes and fraction of CD8 T cells for each subject (*n* = 65 total). Line and shadow indicate linear regression fit and 95% confidence interval, respectively. Significance of the correlations (Pearson *r*) were calculated with a two-sided permutation test. (d-e) Dotplot showing enrichment of pathways (KEGG database) for upregulated genes in HUVECs co-incubated with iMS1 vs. iMono (FDR < 0.1, edgeR exact test). Sizes of circles are proportional to the number of gene hits in a set, whereas color represents the enrichment score of each gene set. (f) Volcano plot showing differential expression analysis results (exact test) between TNFα-activated HUVECs (left) or HREs (right) incubated with iMono or iMS1 conditioned media. Genes with FDR < 0.1 are highlighted in red. In (d-f), *n* = 8 experiments were performed for each condition (2 bone marrow donors with 2 biological and 2 technical replicates)

**Supplementary Figure 10.**
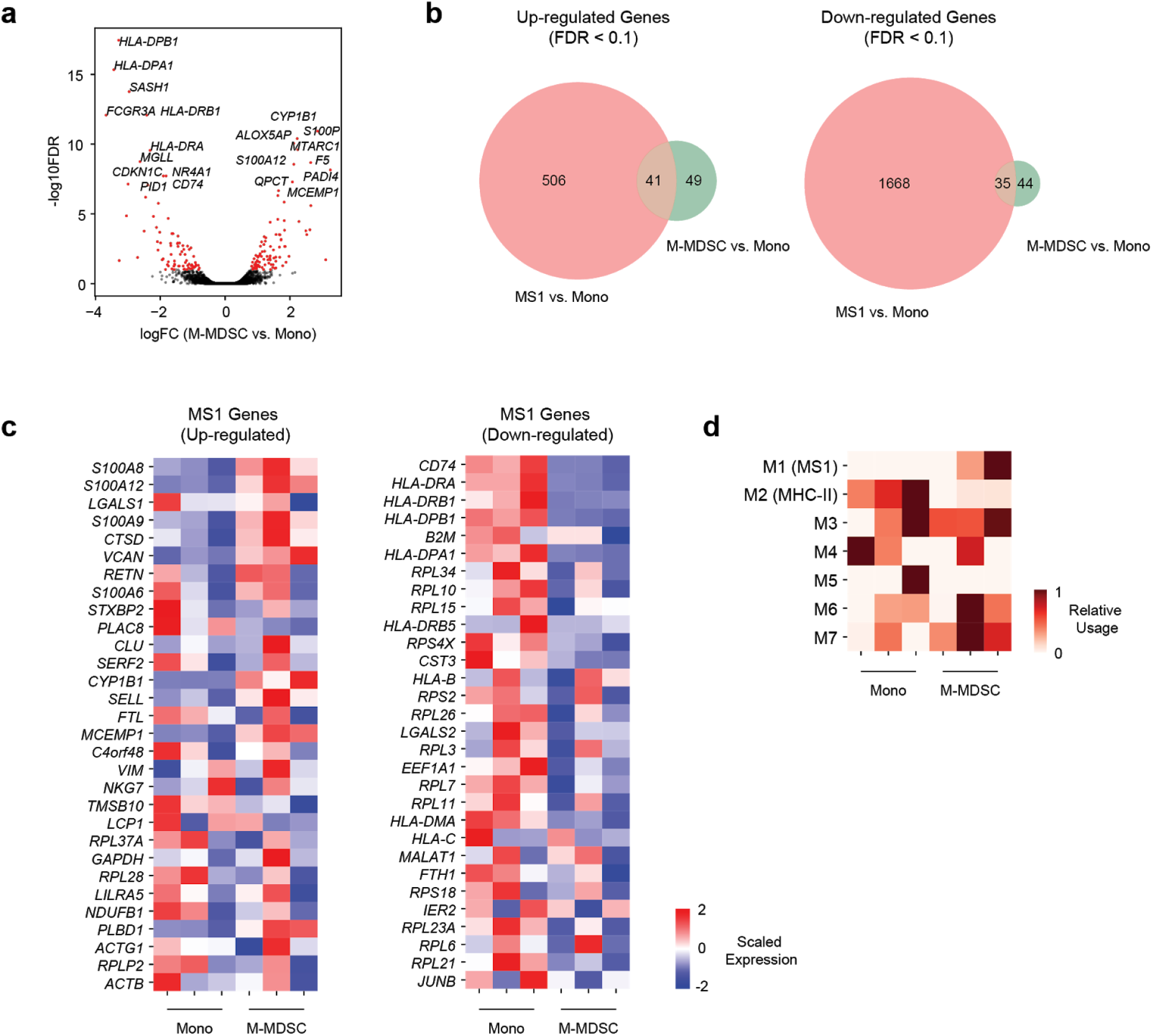
Comparison of MS1 with M-MDSCs. (a) Volcano plot showing differential expression analysis results (exact test) between M-MDSCs and monocytes (*n* = 3 for each) in lung cancer patients from Mastio, *et. al*.^*53*^. Genes with FDR < 0.1 are highlighted in red. (b) Venn diagrams showing overlap of differentially expressed genes (FDR < 0.1, exact test) for MS1 vs. monocytes and M-MDSCs vs. monocytes. (c) Heatmaps showing the expression of the top 30 up-regulated (left) or down-regulated (right) MS1 genes in bulk RNA-seq of monocytes and M-MDSCs (*n* = 3 patients for each). (d) Relative usage of monocyte modules in monocytes and M-MDSCs.

## REFERENCES

1. Rudd, K. E. et al. Global, regional, and national sepsis incidence and mortality, 1990-2017: analysis for the Global Burden of Disease Study. Lancet 395, 200–211 (2020).

2. Filbin, M. R. et al. Presenting Symptoms Independently Predict Mortality in Septic Shock: Importance of a Previously Unmeasured Confounder. Crit. Care Med. 46, 1592–1599 (2018).

3. Scicluna, B. P. & Baillie, J. K. The Search for Efficacious New Therapies in Sepsis Needs to Embrace Heterogeneity. Am. J. Respir. Crit. Care Med. (2018) DOI: 10.1164/rccm.201811-2148ED.

4. van der Poll, T., van de Veerdonk, F. L., Scicluna, B. P. & Netea, M. G. The immunopathology of sepsis and potential therapeutic targets. Nat. Rev. Immunol. 17, 407–420 (2017).

5. Alhazzani, W. et al. Surviving Sepsis Campaign: Guidelines on the Management of Critically Ill Adults with Coronavirus Disease 2019 (COVID-19). Crit. Care Med. 48, e440–e469 (2020).

6. Li, H. et al. SARS-CoV-2 and viral sepsis: observations and hypotheses. Lancet (2020) DOI: 10.1016/S0140-6736(20)30920-X.

7. Arunachalam, P. S. et al. Systems biological assessment of immunity to mild versus severe COVID-19 infection in humans. Science (2020) DOI: 10.1126/science.abc6261.

8. Gupta, A. et al. Extrapulmonary manifestations of COVID-19. Nat. Med. (2020) DOI: 10.1038/s41591-020-0968-3.

9. Reyes, M. et al. An immune-cell signature of bacterial sepsis. Nat. Med. (2020) DOI: 10.1038/s41591-020-0752-4.

10. Drewry, A. M. et al. Persistent lymphopenia after diagnosis of sepsis predicts mortality. Shock 42, 383–391 (2014).

11. Jensen, I. J., Sjaastad, F. V., Griffith, T. S. & Badovinac, V. P. Sepsis-Induced T Cell Immunoparalysis: The Ins and Outs of Impaired T Cell Immunity. J. Immunol. 200, 1543–1553 (2018).

12. Cabrera-Perez, J., Condotta, S. A., Badovinac, V. P. & Griffith, T. S. Impact of sepsis on CD4 T cell immunity. J. Leukoc. Biol. 96, 767–777 (2014).

13. Baudesson de Chanville, C. et al. Sepsis Triggers a Late Expansion of Functionally Impaired Tissue-Vascular Inflammatory Monocytes During Clinical Recovery. Front. Immunol. 11, 675 (2020).

14. Liepelt, A. et al. Differential Gene Expression in Circulating CD14+ Monocytes Indicates the Prognosis of Critically Ill Patients with Sepsis. J. Clin. Med. Res. 9, (2020).

15. Shalova, I. N. et al. Human monocytes undergo functional re-programming during sepsis mediated by hypoxia-inducible factor-1α. Immunity 42, 484–498 (2015).

16. Döcke, W. D. et al. Monocyte deactivation in septic patients: restoration by IFN-gamma treatment. Nat. Med. 3, 678–681 (1997).

17. André, M. C. et al. Bacterial reprogramming of PBMCs impairs monocyte phagocytosis and modulates adaptive T cell responses. J. Leukoc. Biol. 91, 977–989 (2012).

18. Winkler, M. S. et al. Human leucocyte antigen (HLA-DR) gene expression is reduced in sepsis and correlates with impaired TNFα response: A diagnostic tool for immunosuppression? PLoS One 12, e0182427 (2017).

19. Gossez, M. et al. Proof of concept study of mass cytometry in septic shock patients reveals novel immune alterations. Sci. Rep. 8, 17296 (2018).

20. Faivre, V., Lukaszewicz, A.-C. & Payen, D. Downregulation of Blood Monocyte HLA-DR in ICU Patients Is Also Present in Bone Marrow Cells. PLoS One 11, e0164489 (2016).

21. Boomer, J. S. et al. Immunosuppression in patients who die of sepsis and multiple organ failure. JAMA 306, 2594–2605 (2011).

22. Janols, H. et al. A high frequency of MDSCs in sepsis patients, with the granulocytic subtype dominating in gram-positive cases. J. Leukoc. Biol. 96, 685–693 (2014).

23. Hollen, M. K. et al. Myeloid-derived suppressor cell function and epigenetic expression evolves over time after surgical sepsis. Crit. Care 23, 355 (2019).

24. Dai, J., Kumbhare, A., Youssef, D., McCall, C. E. & El Gazzar, M. Intracellular S100A9 Promotes Myeloid-Derived Suppressor Cells during Late Sepsis. Front. Immunol. 8, 1565 (2017).

25. Ulas, T. et al. S100-alarmin-induced innate immune programming protects newborn infants from sepsis. Nat. Immunol. 18, 622–632 (2017).

26. Derive, M., Bouazza, Y., Alauzet, C. & Gibot, S. Myeloid-derived suppressor cells control microbial sepsis. Intensive Care Med. 38, 1040–1049 (2012).

27. Brudecki, L., Ferguson, D. A., McCall, C. E. & El Gazzar, M. Myeloid-derived suppressor cells evolve during sepsis and can enhance or attenuate the systemic inflammatory response. Infect. Immun. 80, 2026–2034 (2012).

28. Schulte-Schrepping, J. et al. Severe COVID-19 is marked by a dysregulated myeloid cell compartment. Cell doi: 10.1016/j.cell.2020.08.001.

29. Kotliar, D. et al. Single-cell profiling of Ebola virus infection in vivo reveals viral and host transcriptional dynamics. bioRxiv 2020.06.12.148957 (2020) DOI: 10.1101/2020.06.12.148957.

30. Kotliar, D. et al. Identifying gene expression programs of cell-type identity and cellular activity with single-cell RNA-Seq. Elife 8, 310599 (2019).

31. Cheng, P. et al. Inhibition of dendritic cell differentiation and accumulation of myeloid-derived suppressor cells in cancer is regulated by S100A9 protein. J. Exp. Med. 205, 2235–2249 (2008).

32. Bah, I., Kumbhare, A., Nguyen, L., McCall, C. E. & El Gazzar, M. IL-10 induces an immune repressor pathway in sepsis by promoting S100A9 nuclear localization and MDSC development. Cell. Immunol. 332, 32–38 (2018).

33. Sweeney, T. E. et al. A community approach to mortality prediction in sepsis via gene expression analysis. Nat. Commun. 9, 694 (2018).

34. Wilk, A. J. et al. A single-cell atlas of the peripheral immune response in patients with severe COVID-19. Nat. Med. 1–7 (2020).

35. Lee, J. S. et al. Immunophenotyping of COVID-19 and influenza highlights the role of type I interferons in development of severe COVID-19. Sci Immunol 5, (2020).

36. Veglia, F., Perego, M. & Gabrilovich, D. Myeloid-derived suppressor cells coming of age. Nat. Immunol. 19, 108–119 (2018).

37. Mehta, P. et al. COVID-19: consider cytokine storm syndromes and immunosuppression. Lancet 395, 1033 (2020).

38. Wu, L. et al. Ascites-derived IL-6 and IL-10 synergistically expand CD14+HLA-DR-/low myeloid-derived suppressor cells in ovarian cancer patients. Oncotarget 8, 76843–76856 (2017).

39. Mundy-Bosse, B. L. et al. Distinct myeloid suppressor cell subsets correlate with plasma IL-6 and IL-10 and reduced interferon-alpha signaling in CD4 ^+^ T cells from patients with GI malignancy. Cancer Immunol. Immunother. 60, 1269–1279 (2011).

40. Braun, D. A., Fribourg, M. & Sealfon, S. C. Cytokine response is determined by duration of receptor and signal transducers and activators of transcription 3 (STAT3) activation. J. Biol. Chem. 288, 2986–2993 (2013).

41. Niemand, C. et al. Activation of STAT3 by IL-6 and IL-10 in primary human macrophages is differentially modulated by suppressor of cytokine signaling 3. J. Immunol. 170, 3263–3272 (2003).

42. Stec, M. et al. Expansion and differentiation of CD14+CD16(-) and CD14+ +CD16+ human monocyte subsets from cord blood CD34+ hematopoietic progenitors. J. Leukoc. Biol. 82, 594–602 (2007).

43. Marigo, I. et al. Tumor-induced tolerance and immune suppression depend on the C/EBPbeta transcription factor. Immunity 32, 790–802 (2010).

44. Uhel, F. et al. Early Expansion of Circulating Granulocytic Myeloid-derived Suppressor Cells Predicts Development of Nosocomial Infections in Patients with Sepsis. Am. J. Respir. Crit. Care Med. 196, 315–327 (2017).

45. Lelubre, C. & Vincent, J.-L. Mechanisms and treatment of organ failure in sepsis. Nat. Rev. Nephrol. 14, 417–427 (2018).

46. Martin, G. et al. Role of plasma matrix-metalloproteases (MMPs) and their polymorphisms (SNPs) in sepsis development and outcome in ICU patients. Sci. Rep. 4, 5002 (2014).

47. Iba, T. et al. Diagnosis and management of sepsis-induced coagulopathy and disseminated intravascular coagulation. J. Thromb. Haemost. 17, 1989–1994 (2019).

48. Dhainaut, J.-F. et al. Dynamic evolution of coagulopathy in the first day of severe sepsis: relationship with mortality and organ failure. Critical care medicine vol. 33 341–348 (2005).

49. Merad, M. & Martin, J. C. Pathological inflammation in patients with COVID-19: a key role for monocytes and macrophages. Nat. Rev. Immunol. 20, 355–362 (2020).

50. Ramlall, V. et al. Immune complement and coagulation dysfunction in adverse outcomes of SARS-CoV-2 infection. Nat. Med. (2020) DOI: 10.1038/s41591-020-1021-2.

51. Harrison, C. Focus shifts to antibody cocktails for COVID-19 cytokine storm. Nat. Biotechnol. 38, 905–908 (2020).

52. Garbers, C., Heink, S., Korn, T. & Rose-John, S. Interleukin-6: designing specific therapeutics for a complex cytokine. Nat. Rev. Drug Discov. 17, 395–412 (2018).

53. Mastio, J. et al. Identification of monocyte-like precursors of granulocytes in cancer as a mechanism for accumulation of PMN-MDSCs. J. Exp. Med. 216, 2150–2169 (2019).

54. Höchst, B. et al. Activated human hepatic stellate cells induce myeloid derived suppressor cells from peripheral blood monocytes in a CD44-dependent fashion. J. Hepatol. 59, 528–535 (2013).

55. Solito, S. et al. A human promyelocytic-like population is responsible for the immune suppression mediated by myeloid-derived suppressor cells. Blood 118, 2254–2265 (2011).

56. Heine, A. et al. Generation and functional characterization of MDSC-like cells. Oncoimmunology 6, e1295203 (2017).

57. Stoeckius, M. et al. Cell Hashing with barcoded antibodies enables multiplexing and doublet detection for single cell genomics. Genome Biol. 19, 224 (2018).

58. Villani, A.-C. et al. Single-cell RNA-seq reveals new types of human blood dendritic cells, monocytes, and progenitors. Science 356, eaah4573–eaah4573 (2017).

59. Wolf, F. A., Angerer, P. & Theis, F. J. SCANPY: large-scale single-cell gene expression data analysis. Genome Biol. 19, 15 (2018).

60. Bergen, V., Lange, M., Peidli, S., Wolf, F. A. & Theis, F. J. Generalizing RNA velocity to transient cell states through dynamical modeling. Nat. Biotechnol. (2020) DOI: 10.1038/s41587-020-0591-3.

61. Newman, A. M. et al. Robust enumeration of cell subsets from tissue expression profiles. Nat. Methods 12, 453–457 (2015).

62. Haynes, W. A. et al. Empowering Multi-Cohort Gene Expression Analysis to Increase Reproducibility. In Biocomputing 2017 144–153 (World Scientific, 2016).

63. Nakahira, K. et al. Circulating mitochondrial DNA in patients in the ICU as a marker of mortality: derivation and validation. PLoS Med. 10, e1001577. discussion e1001577 (2013).

64. Dolinay, T. et al. Inflammasome-regulated cytokines are critical mediators of acute lung injury. Am. J. Respir. Crit. Care Med. 185, 1225–1234 (2012).

65. Picelli, S. et al. Full-length RNA-seq from single cells using Smart-seq2. Nat. Protoc. 9, 171–181 (2014).

66. Reyes, M. et al. Multiplexed enrichment and genomic profiling of peripheral blood cells reveal subset-specific immune signatures. Science Advances 5, eaau9223 (2019).

67. Solito, S. et al. Methods to Measure MDSC Immune Suppressive Activity In Vitro and In Vivo. Curr. Protoc. Immunol. 124, e61 (2019).

